# Macaque Vγ4⁺ γδ T cells exhibit distinct IFN-γ and IL-22 effector profiles that are dysregulated during chronic SIV infection and diarrheal disease

**DOI:** 10.64898/2026.11.13.766599

**Authors:** Alexandra B McNally, Andrew R Rahmberg, Tovah E. Markowitz, Victoria A Hart, Jesse Do, Siva Thirugnanam, Yoshinori Fukazawa, Nizar Batada, Jason M Brenchley, Namita Rout

## Abstract

Vγ4⁺ γδ T cells contribute to colonic homeostasis in humans, but their biology in nonhuman primates remains poorly defined. Here, we characterized Vγ4⁺ γδ T cells in blood and intestinal tissues of healthy, diarrhea-inflicted, and SIV-infected rhesus macaques. TRG/TRD repertoire sequencing across colon, mesenteric lymph node, blood, and spleen revealed compartment-specific clonal architecture and colonic enrichment of TRGV3. Flow cytometric profiling revealed circulating and colonic Vγ4⁺ T cells were predominantly Vδ1⁺, exhibited an effector memory phenotype, and displayed T-bet-dominant transcriptional profiles. High-dimensional clustering confirmed Vγ4⁺Vδ1⁺ T cells as a distinct terminally differentiated lineage. Functionally, stimulated Vγ4⁺ T cells exhibited robust Th1/Th17 cytokine production, supporting dual inflammatory and barrier-protective roles. Campylobacter-associated diarrhea was characterized by mucosal expansion of central memory Vγ4⁺ T cells and altered cytokine responses. In contrast, ART-suppressed chronic SIV infection was associated with colonic enrichment of naïve-like Vγ4⁺ T cells and increased RORγt expression, alongside profound systemic loss of cytokine production but relative preservation of mucosal Th17-type responses. Both conditions exhibited a convergent loss of systemic Type 1 effector Vγ4⁺ T-cell responses, pointing to a shared functional alteration during chronic intestinal inflammation. Overall, these findings reveal disease-specific remodeling of Vγ4⁺ T cells during chronic intestinal inflammation and gut barrier disruption in chronic enterocolitis and SIV infection.

**Author Summary:** γδ T cells contribute to intestinal barrier immunity by balancing antimicrobial defense and tissue homeostasis. In humans, the subset of Vγ4⁺ γδ T cells has been implicated in maintaining colonic homeostasis, but their phenotypic and functional characteristics in nonhuman primates remain unexplored. In this study, we provide the first comprehensive characterization of rhesus macaque Vγ4⁺ γδ T cells under healthy conditions and during Campylobacter-associated diarrhea, and ART-suppressed SIV infection, as two distinct settings of pathogen-driven chronic intestinal inflammation. We show that these cells represent a distinct, predominantly Vδ1⁺ effector-memory population capable of producing both type 1 and type 3 cytokines, suggesting roles in antimicrobial defense and mucosal immunity. Although both disease conditions were associated with impaired systemic type 1 effector function, their effects on intestinal Vγ4⁺ γδ T cells differed. Campylobacter-associated diarrhea was characterized by mucosal expansion of central memory cells, whereas chronic SIV infection was associated with enrichment of naïve-like cells and increased RORγt expression, with relative preservation of colonic type 3 responses. Together, these findings highlight disease-specific mucosal remodeling and shared systemic functional impairment of Vγ4⁺ γδ T cells, providing new insights into their contribution to pathogen defense and intestinal barrier functions during chronic gut inflammation.

## Introduction

Gamma delta (γδ) T cells are unconventional T lymphocytes that bridge innate and adaptive immunity and are strategically positioned at mucosal and epithelial barrier surfaces(1). Although they comprise only a small fraction of circulating T cells in humans, γδ T cells are highly enriched within barrier tissues, where they play central roles in maintaining tissue homeostasis, coordinating antimicrobial defense, promoting epithelial repair, and regulating inflammatory responses(2). Beyond their well-established contributions to antiviral and antitumor immunity(3–5), γδ T cells are increasingly recognized as key orchestrators of mucosal integrity and immune surveillance(6), enabling rapid responses to tissue stress, infection, and injury.

Despite their importance, the biology of human γδ T cells remains incompletely understood, in part because they differ substantially from their murine counterparts in TCR repertoire usage, antigen-recognition mechanisms, tissue distribution, and functional specialization(7). Like conventional αβ T cells, γδ T cells express clonotypic T cell receptors generated through somatic recombination; however, γδ TCRs recognize a distinct spectrum of ligands and stress-induced signals(8), allowing these cells to integrate tissue-derived cues with immune effector functions. Defining the tissue-specific composition and roles of human γδ T cell subsets therefore remains a central challenge in mucosal immunology. Macaque γδ T cells are remarkably similar to human γδ T cells, sharing up to 91% amino acid sequence homology in their receptor elements(9–11). This close evolutionary relationship makes macaques the primary preclinical model for studying human immune biology.

Human γδ T cells comprise two major subsets identified by their Vδ chain usage: Vδ1⁺ T cells that are predominant in the thymus and peripheral tissues and recognize stress-related antigens(12, 13), and Vδ2⁺ T cells that constitute most blood γδ T cells and predominantly recognize phosphorylated non-peptidic antigens that are metabolic intermediates of the isoprenoid biosynthesis(14, 15). However, γδ T cell function is also influenced by pairing with specific Vγ chains, particularly in barrier tissues where butyrophilin (BTN/BTNL)-mediated selection and local antigenic cues shape tissue-specific γδ T-cell compartments(16, 17). Recent studies have identified human Vγ4⁺ T cells as a distinct BTNL3/BTNL8-responsive population within the colon, where they regulate epithelial homeostasis and are implicated in the pathogenesis of inflammatory bowel diseases (IBDs), including Crohn’s disease (CD) and ulcerative colitis (UC)(18–20). Despite growing evidence linking Vγ4⁺ T cells to intestinal health, their biology outside the context of IBD remains poorly understood, and corresponding populations have not been defined in nonhuman primates.

Here, we identify and characterize the rhesus macaque counterpart of human Vγ4⁺ T cells, defining their structural, phenotypic, and functional properties in peripheral blood and colonic mucosa. We further demonstrate how the intestinal Vγ4⁺ T cell compartment is differentially remodeled under two distinct models of chronic intestinal disease, *Campylobacter*-driven enterocolitis and chronic SIV infection. These findings advance our understanding of γδ T cell-mediated barrier immunity and reveal potential mechanisms underlying mucosal immune failure in chronic infectious disease.

## Results

### γδTCR chain usage and clonal space across systemic and mucosal compartments

To define systemic versus tissue-resident γδ T cell clonal architecture, we sequenced PCR-amplified TRG and TRD V–J junctions from sorted αβTCR-negative T cells across the colon, mesenteric lymph nodes (MLN), PBMC, and spleen of rhesus macaques. After filtering unique sequences to minimize errors, clonal diversity was assessed by frequency group abundance. The TRG repertoire was dominated by large-frequency clonotypes, whereas the TRD repertoire contained more hyperexpanded clonotypes, reflecting lower overall TRD diversity (Fig 1A, B). Aggregated by tissue, the spleen exhibited higher proportions of medium-frequency TRG and TRD clonotypes, indicating greater compartmental diversity (Fig 1C, D).

**Fig 1.**
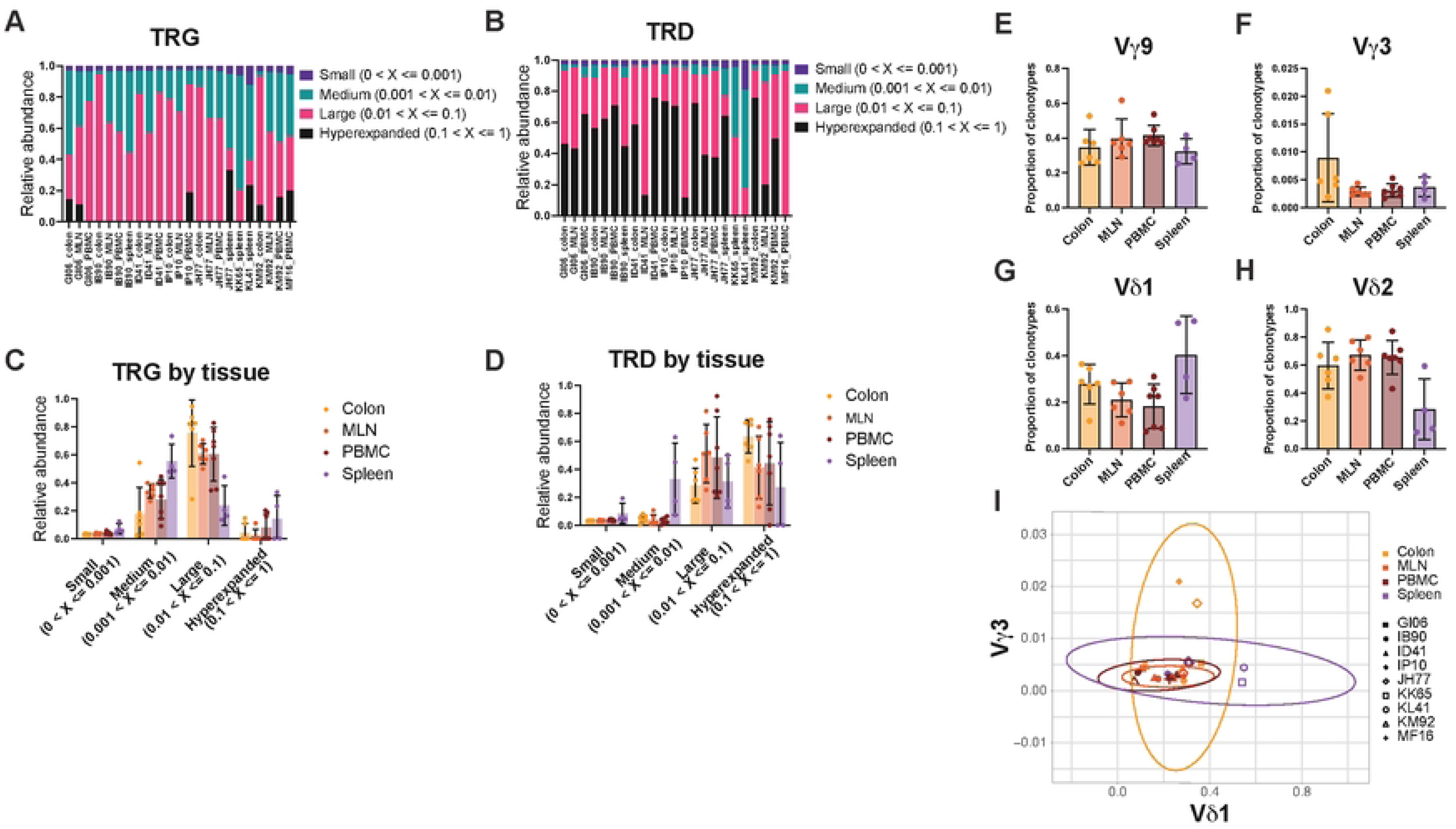
Characterization of the Rhesus Macaque γδ TCR Repertoire across Systemic and Mucosal Microenvironments. High-throughput T cell receptor (TCR) sequencing was performed on genomic DNA isolated from αβTCR^−^ T cells from the colon, mesenteric lymph nodes (MLN), peripheral blood mononuclear cells (PBMC), and spleen tissue (n=4-6 rhesus macaques). **(A-B)** Representative individual stacked bar profiles demonstrating the occupied repertoire space of the **(A)** T cell receptor γ (TRG) and **(B)** T cell receptor δ (TRD) chains across individual animals, stratified by clonotype indices (clonotype abundance bins ranging from unique/low [2:5] to highly expanded [301:10000]). **(C-D)** Combined analysis of occupied repertoire space grouped by anatomical tissue compartment for **(C)** TRG and **(D)** TRD. Bars indicate the mean proportion of the total repertoire space occupied by clones within each designated abundance bin SEM. **(E-H)** Proportions of total detected clonotypes utilizing specific variable region segments, including **(E)** Vδ2, **(F)** Vγ9, **(G)** Vδ1, and **(H)** Vγ3. Each data point represents an individual animal; error bars indicate mean SEM. Note that rhesus *TRGV3* represents the closest genetic sequence homolog to human *TRGV4*, identifying it as the human lineage equivalent via BLAST alignment. Statistical analysis via one-way ANOVA with multiple comparisons revealed that colonic enrichment of the subset approaches statistical significance against systemic tissues (colon vs. MLN, P=0.0540; colon vs. PBMC, P=0.0537). **(I)** Clustering analysis mapping individual tissue samples-based frequency of Vd1 and Vg3 usage. Shaded areas represent the 95% confidence intervals (ellipses) for each tissue compartment. Overlapping ellipses demonstrate a shared structural framework, though the colon population demonstrates a vertical elongation driven by higher-than-average and co-utilization. Individual animal identifiers are indicated by corresponding geometric symbols in the legend.

To determine whether tissue-specific clonal distributions reflected distinct structural γδ T-cell subsets, we quantified variable (V) gene usage across compartments. PBMC and gut-draining MLN displayed coordinated TCR profiles dominated by Vδ2 and Vγ9 chains, comprising 40–60% of total clonotypes (Fig 1E, F). Systemic Vδ1 usage varied inter-individually, showing spleen enrichment in two of four animals (Fig 1G). Notably, TRGV3 was rare across PBMC, MLN, and spleen (<0.2%) but selectively enriched in the colonic lamina propria (1–2% of local TRG repertoire; p=0.0540 vs. MLN, p=0.0537 vs. PBMC; Fig 1H), suggesting tissue-associated specialization of this clonotype/variable-gene family.

To evaluate potential pairing dynamics within this mucosal niche, we correlated TRGV3 and TRDV1 frequencies from matching tissue samples. The two colonic samples exhibiting the highest overall TRGV3 usage also clustered above the mean for TRDV1. Overall, while the majority of colon samples exhibited a dual trend toward higher-than-average usage of both TRGV3 and TRDV1, this shift was not pronounced enough to drive a complete spatial separation. As a result, the 95% confidence intervals for the colon and systemic compartments overlap within the ellipse visualization (Fig1 I). These findings therefore support a colon-associated shift in TRGV3 and TRDV1 usage within the broader γδ TCR repertoire, rather than a discrete or exclusive TRGV3/TRDV1-defined population. Importantly, the repertoire analysis does not establish evolutionary equivalence between rhesus TRGV3 and any specific human TRGV lineage, nor does TRGV3 usage establish the molecular identity of the population detected by the anti-human Vγ4 antibody. We therefore considered TRGV3 enrichment as an independent observation of rhesus γδ TCR repertoire structure and did not use it to define the antibody-detected population. To characterize the phenotype and effector programs of the γδ T-cell population detected by the anti-human Vγ4 antibody, we next used high-parameter flow cytometry.

### Circulating macaque Vγ4^+^ T cells predominantly utilize the Vδ1 chain and exhibit an effector memory phenotype

To characterize steady-state Vγ4^+^ T cells, we evaluated Vδ1^+^, Vδ2^+^, and Vγ4^+^ T cell phenotypes across healthy rhesus macaque blood, liver, and colon lamina propria (Fig 2A). In contrast to Vδ2^+^ T cells, which were enriched in PBMCs and liver (P < 0.0001), Vγ4^+^ T cells maintained low, uniform frequencies across compartments (Fig 2B). Vγ4^+^ T cell frequencies strongly correlated with Vδ1^+^ subset in both PBMCs (r = 0.83, P < 0.0001; Fig 2C) and colonic LPLs (r = 0.8571, P = 0.0238; Supplementary Fig 1A), with most co-expressing Vδ1. Conversely, Vγ4^+^ and Vδ2^+^ T cell frequencies were uncoupled in blood (r = –0.35, P = 0.096; Fig 2D) and colon (r = 0, P > 0.99; Supplementary Fig 1B). Given the role of human Vγ4^+^Vδ1^+^ T cells in mucosal homeostasis and inflammation(18, 20, 21), we prioritized blood and colon for downstream profiling.

**Fig 2.**
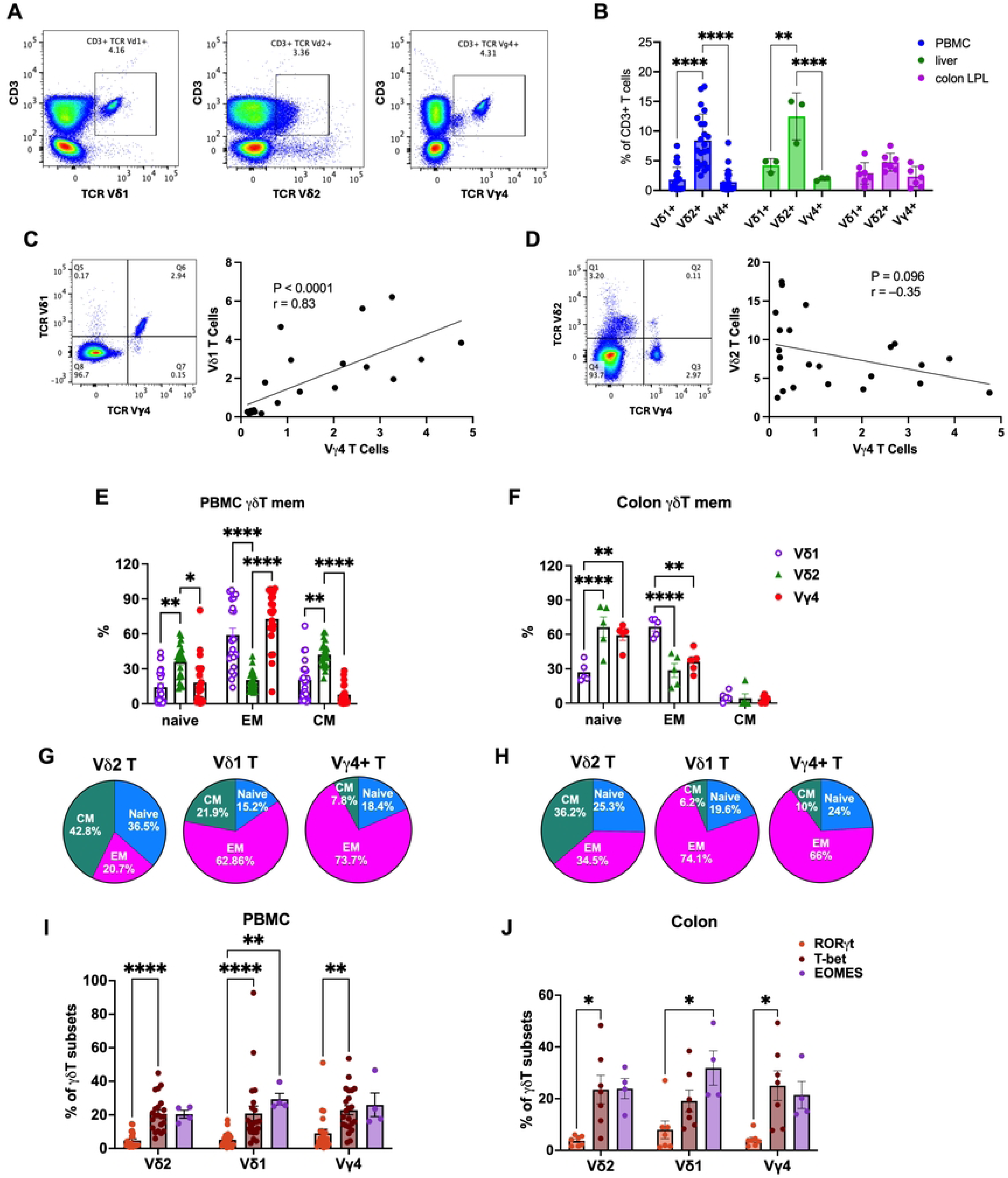
Phenotypic characterization of macaque Vγ4^+^ T cells. **(A)** Representative flow cytometry plots showing gating of Vδ1^+^, Vδ2^+^, and Vγ4^+^ subsets within live CD3^+^ lymphocytes from healthy rhesus macaque PBMCs. **(B)** Frequency of Vδ1^+^, Vδ2^+^, and Vγ4^+^ T cell subsets as a percentage of CD3^+^ T cells in PBMCs, liver, and colon LPL from healthy animals (PBMC, n=22; colon, n=7; liver, n=3). Statistical comparisons were performed by two-way ANOVA with Tukey’s multiple comparisons correction. **(C)** Representative co-expression dot plot and correlation between Vγ4 and Vδ1 frequency in PBMCs (n=22) showing strong positive correlation (Spearman r=0.83, p<0.0001). **(D)** Representative co-expression dot plot and correlation between Vγ4 and Vδ2 frequency in PBMCs (n=22). **(E-F)** Frequency of naive, effector memory (EM), and central memory (CM) phenotypes within Vδ1^+^, Vδ2^+^, and Vγ4^+^ populations in PBMCs and colon LPL from healthy animals (PBMC, n=22; colon LPL, n=7). **(G-H)** Pie charts depicting mean memory composition of Vδ2^+^, Vδ1^+^, and Vγ4^+^ in PBMCs **(G)** and colon LPL **(H)**. **(I-J)** Expression of transcription factors RORγt, T-bet, and EOMES as a percentage of each γδ T subset in PBMCs **(I),** and colon LPL **(J)** from healthy animals (n=7–22). Bars represent mean ± SEM. Statistical comparisons were performed by two-way ANOVA with Tukey’s multiple comparisons correction. *p<0.05, **p<0.01, ***p<0.001, ****p<0.0001.

We next evaluated differentiation profiles using CD28 and CD95 to define naïve (CD28^+^CD95^−^), central memory (T_CM_; CD28^+^CD95^+^), and effector memory (T_EM_; CD28^−^CD95^+^) subsets (Fig 2E, F). Both Vδ1^+^ and Vγ4^+^ T cells were heavily skewed toward a T_EM_ phenotype (P < 0.0001 vs. Vδ2^+^; Fig 2E). In contrast, Vδ2^+^ T cells were characterized by higher naïve cell frequencies (P < 0.05 vs. Vδ1^+^and Vγ4^+^) and prominent T_CM_ representation, which significantly exceeded both the Vδ1^+^and Vγ4^+^ pools (P < 0.01 Vδ1^+^and P < 0.001 vs. Vγ4^+^). This pattern was largely mirrored in the colon lamina propria, where Vδ1^+^ cells and Vγ4^+^ cells remained highly T_EM_-skewed (P < 0.007 vs. Vδ2^+^; Fig 2F), while Vδ2^+^ T cells maintained significantly higher T_CM_ frequencies than Vδ1^+^ (P = 0.0003) and Vγ4^+^ cells (P = 0.0024). Compositional pie chart analysis highlighted the T_EM_-dominant phenotype of the Vγ4^+^ population similar to Vδ1^+^ T cells (Fig. 1G, H) in both PBMCs (73.74%; Fig 2G) and colon LPL (66.09%; Fig 2H), with sparse T_CM_ representation (≤10%). Conversely, Vδ2^+^ T cells preserved robust T_CM_ fractions across both systemic and mucosal compartments (42.78% in PBMCs, 36.25% in colon LPL). Collectively, these data demonstrate that while macaque Vδ2^+^ T cells retain a balanced memory architecture, Vγ4^+^ T cells exist predominantly as terminally differentiated effector cells primed for immunosurveillance at mucosal barriers.

Flow cytometric quantification of lineage-defining transcription factors (RORγt, T-bet, and EOMES) revealed greater T-bet and EOMES expression in contrast to minimal RORγt expression by all three subsets displayed in peripheral blood (Fig 2I). Notably, circulating γδ T cells exhibited significantly higher T-bet frequencies compared to RORγt (P < 0.0001 for Vδ2^+^ and Vδ1^+^; and P < 0.01 for Vγ4^+^), while the Vδ1^+^ subset was characterized by a distinct enrichment of EOMES as well (P < 0.01). This Th1 type/cytolytic potential was maintained within the colon lamina propria, where T-bet or EOMES expression remained significantly elevated over RORγt across all subsets (P < 0.05; Fig 2J). Thus, under steady state conditions, macaque Vγ4^+^ T cells share a dominant cytolytic and Type 1 effector program with both the Vδ1^+^and Vδ2^+^ subsets in peripheral blood and colon tissue.

To map the Vγ4^+^ T cells within the broader landscape of conventional T cells and γδ T cells, we performed unsupervised FlowSOM clustering and t-SNE dimensionality reduction on total CD45^+^ live lymphocytes from PBMCs (Fig 3A-C). Automated metaclustering identified eight distinct cellular populations based on hierarchical marker expression topologies (Fig 3B). High-dimensional t-SNE projection revealed that Vγ4^+^Vδ1^+^ T cells clustered into a unique, highly cohesive island (population 5) that was spatially segregated from conventional CD4^+^ and CD8^+^ T-cell pools. Hierarchical clustering heat-map confirmed that this Vγ4^+^Vδ1^+^ T cell cluster was defined by coordinated, high-level expression of both TCR Vγ4 and TCR Vδ1 in a CD4-negative configuration, paired with a T-bet^+^CD95^+^CD28^−^ effector memory profile (Fig 3B). In sharp contrast, Vδ2^+^ T cells partitioned into a structurally distinct cluster (population 4) characterized by prominent CD28 expression, reflecting a less-differentiated memory profile. Finally, minimal spanning tree topology demonstrated that while the Vδ2^+^ lineage branched independently adjacent to the CD4^+^ T-cell trajectory, the Vγ4^+^Vδ1^+^ cluster anchored as a highly differentiated terminal branch extending directly from the CD8^+^ effector memory arm (Fig 3C), reinforcing their highly specialized cytolytic polarization.

**Fig 3.**
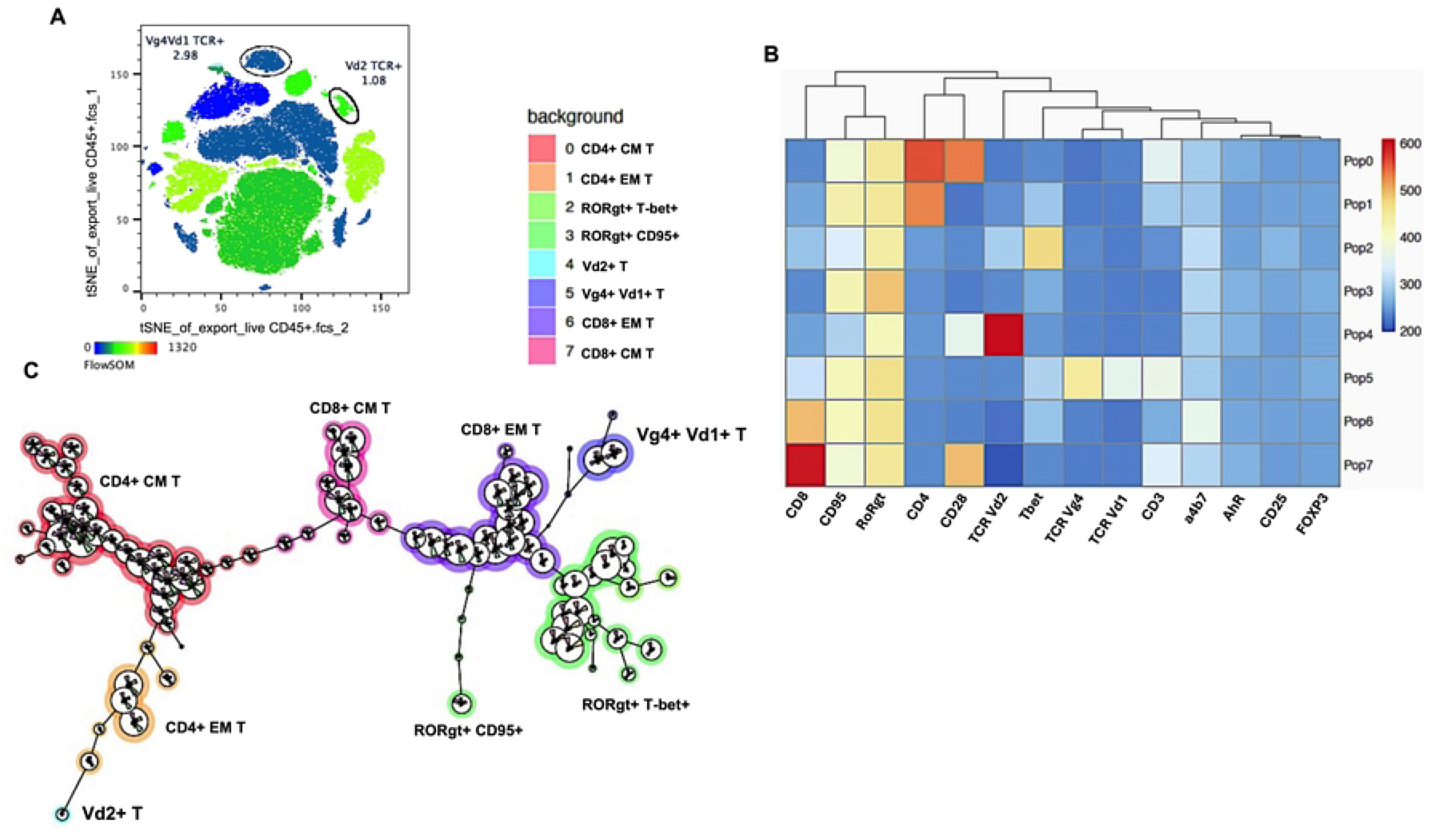
High-dimensional unsupervised clustering defines a distinct effector trajectory for circulating Vγ4^+^ Vδ1^+^ T cells in rhesus macaques. **(A)** Unsupervised t-distributed Stochastic Neighbor Embedding (t-SNE) map of total live CD45^+^ lymphocytes from healthy rhesus macaque PBMCs. **(B)** Hierarchical clustering and heatmap displaying the median fluorescence intensity (MFI) of 15 lineage-defining surface markers and transcription factors across the eight automated FlowSOM metaclusters (Pop0 to Pop7). The dendrogram (top) establishes phenotypic relatedness based on global multidimensional marker expression topologies. Color scale ranges from low (blue) to high (red) relative expression intensity. **(C)** Minimal Spanning Tree visualization of the FlowSOM clustering hierarchy. Individual nodes represent localized single-cell phenotypic clusters, connected by branches based on short multi-dimensional distance vectors. Background color shading corresponds to the designated leukocyte lineages tracked in the t-SNE background panel, illustrating the structural divergence of the Vδ2^+^ trajectory from the terminal Vγ4^+^Vδ1^+^ T branch extending from the CD8^+^ T_EM_ arm. Data are representative of 3 healthy rhesus macaques.

### Vγ4^+^ T cells display predominant IFN-γ and IL-22 cytokine functions

To evaluate the functional capacity and polarization of macaque Vγ4^+^ T-cell subsets, we profiled intracellular cytokine production following either polyclonal mitogen (PMA/Ionomycin) or plate-bound TCR stimulation (Fig 4). Upon maximal mitogen activation, Vγ4^+^ T cells exhibited a higher cytokine-expression profile in comparison to Vδ1^+^ T cells (Fig 4A). Specifically, Vγ4^+^ T cells demonstrated robust Type 1 and Type 3 effector polarization; mean IFN-γ = 75.54% (P = 0.013 vs. Vδ1^+^) and mean IL-22 = 63.4% (P = 0.043 vs. Vδ1^+^). Furthermore, Vγ4^+^ T cells produced significantly higher levels of TNF-α (P = 0.0009 vs. Vδ1^+^). While Vδ2^+^ T cells displayed a trend of lower IFN-γ and IL-22 production in comparison to Vγ4^+^ T cells, it did not reach statistical significance.

**Fig 4.**
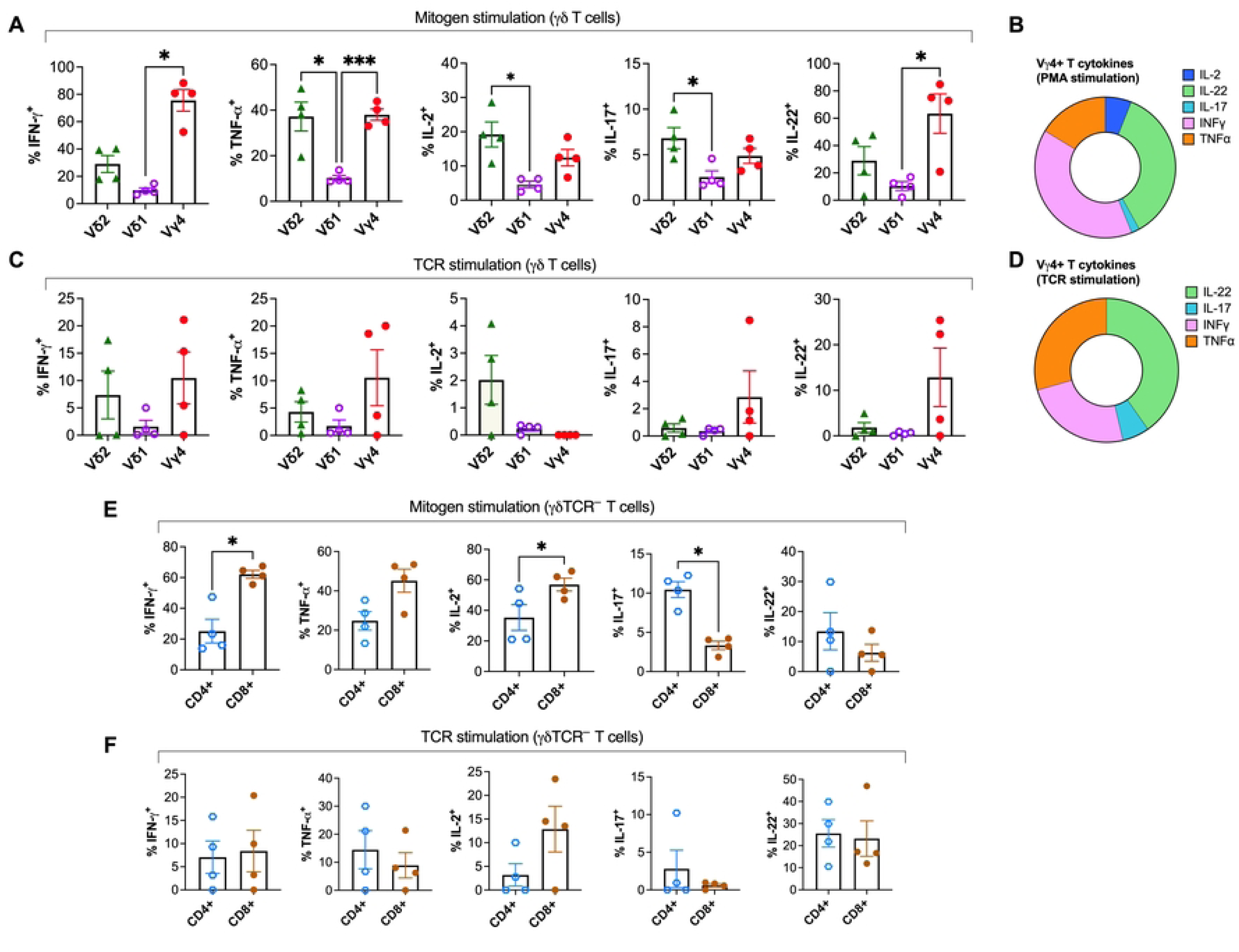
Cytokine-producing profiles of macaque γδ T-cell subsets following mitogen or TCR-specific stimulation. Intracellular cytokine production of IFN-γ, TNF-α, IL-2, IL-17, and IL-22 was quantified by flow cytometry and expressed as a percentage of each γδ T subset in PBMCs from healthy animals. Cells were evaluated following either polyclonal mitogen activation with PMA/ionomycin **(A-B)** or plate-bound TCR crosslinking stimulation **(C-D)**. **(B, D)** Donut charts illustrate the mean relative composition of the total cytokine response within the Vγ4^+^ T cell subset. Bar graphs display individual animal frequencies for each cytokine for γδ T-cell subsets **(A, C),** or TCR γδ-negative CD3^+^ T cell subsets **(E, F)**. Background cytokine production from unstimulated controls was subtracted from all stimulated experimental values; any resulting negative values were set to zero. Bars represent mean ± SEM. Statistical significance was determined using a two-way ANOVA followed by Tukey’s post-hoc multiple comparisons test. *p<0.05, ***p<0.001.

Qualitative compositional analysis via donut charts confirmed this unique functional profile, revealing that the Vγ4^+^ T-cytokine response is balanced between classical inflammatory mediators (IFN-γ and TNF-α) and mucosal barrier-protective cytokines (IL-22 and IL-17) (Fig 4B). Notably, this polyfunctional program was preserved under physiological conditions following direct TCR crosslinking (Fig 4C, D) with CD3 and CD28 antibodies. Although absolute frequencies were expectedly lower than those induced by mitogen stimulation, Vγ4^+^ T cells consistently maintained better coordinated expression of IFN-γ/TNF-α, and IL-22) through the TCR complex compared to both the Vδ1^+^ and Vδ2^+^ subsets. Intracellular cytokine staining further showed robust IL-22 production by Vγ4^+^ T cells relative to classical CD4^+^ and CD8^+^ T cells following mitogen stimulation (Fig 4E), albeit this distinction was lost in response to TCR stimulation (Fig 4F). Collectively, these data demonstrate that macaque Vγ4^+^ T cells represent a highly polyfunctional effector lineage uniquely capable of mediating both Th1-type cytokine functions and mucosal barrier defense.

### Intestinal inflammation during chronic diarrheal disease drives Vγ4^+^ and Vδ1^+^ T cell remodeling and RORγt upregulation

To investigate how chronic intestinal inflammation shapes γδ T cell dynamics, we evaluated Vδ1⁺, Vδ2⁺, and Vγ4⁺ T cell frequencies in blood and colonic LPLs during *Campylobacter*-associated chronic gastrointestinal disease (Fig 5A, Table 1). Compared to controls, circulating Vδ1^+^ T cells were significantly greater during chronic diarrhea (P < 0.0001). Similarly, colonic Vγ4^+^ cells demonstrated an upward trend (P = 0.08), whereas Vδ2^+^ subpopulations remained unaltered. This coordinated multi-compartmental shift mirrored steady-state Vγ4/Vδ1 pairing (Fig 2), indicating that this anatomical axis responds dynamically to intestinal inflammation (Fig 5A).

**Fig 5.**
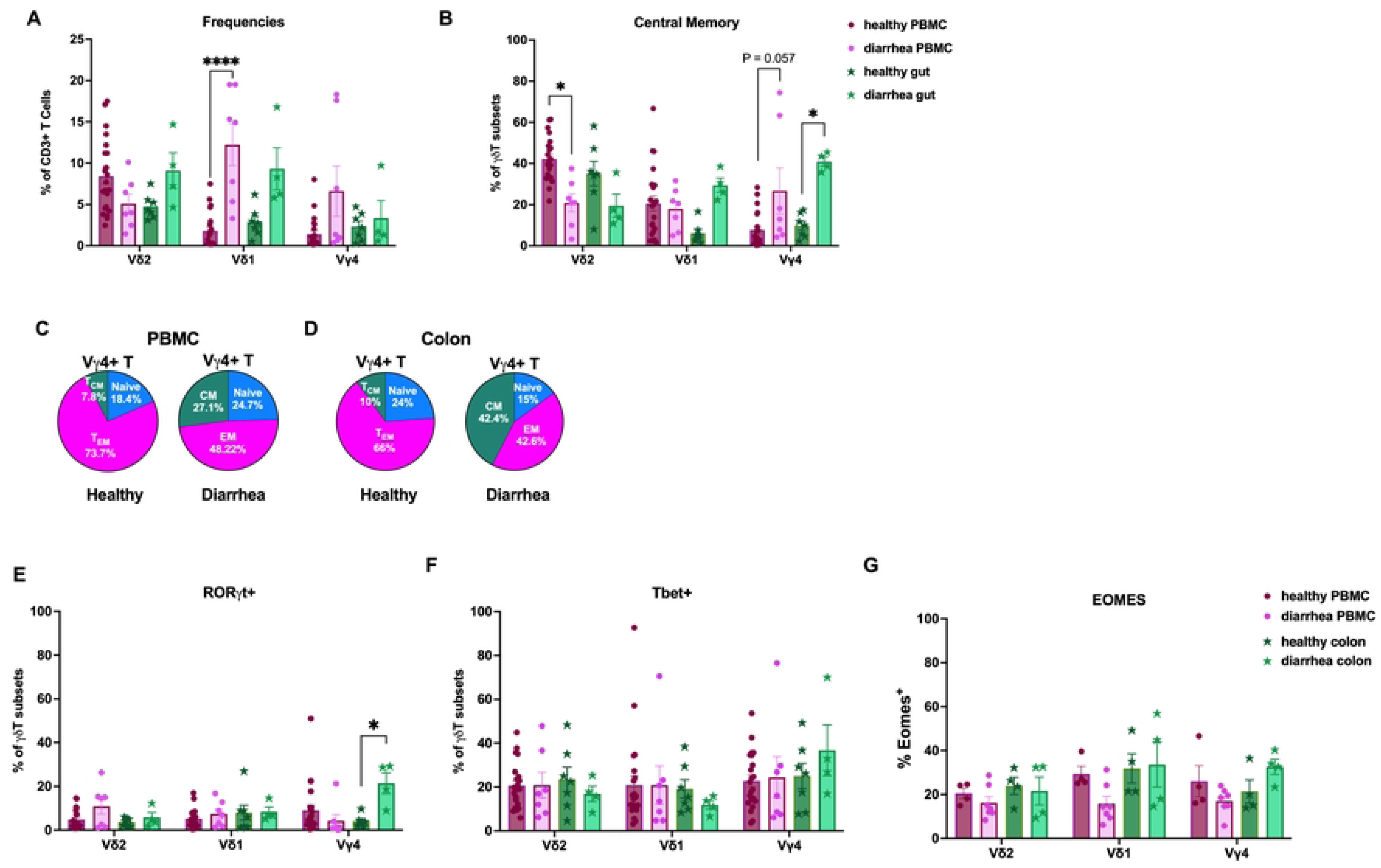
Frequencies and functional phenotype of macaque γδ T-cell subsets during chronic diarrheal disease. **(A)** Frequency of Vδ2, Vδ1, and Vγ4 subsets as a percentage of CD3^+^ T cells in PBMCs and colon LPL from animals with chronic diarrheal disease (PBMC, n=7; colon, n=4) compared to healthy animals (PBMC, n=22; colon, n=7). **(B)** Frequency of CD95^+^CD28^+^ central memory (T_CM_) phenotype cells within Vδ2, Vδ1, and Vγ4 populations in PBMCs and colon LPL from animals with chronic diarrheal disease (PBMC, n=7; colon LPL, n=4) compared to healthy animals (PBMC, n=22; colon LPL, n=7). **(C-D)** Pie charts depicting mean memory composition of Vγ4^+^ subsets in PBMCs **(C)** and colon LPL **(D)** during chronic diarrheal disease (right) compared to steady state animals (left). RORγt **(E)**, T-bet **(F),** and EOMES **(G)** expression as a percentage of each γδ subset in PBMCs and colon LPL in healthy versus chronic diarrheal disease. Bars represent mean ± SEM. Statistical comparisons were performed by two-way ANOVA with Tukey’s multiple comparisons correction. *p<0.05, ****p<0.0001.

**Table 1.** Animal Demographics and Clinical Characteristics.

| **Animal ID** | **Sex** | **Age (yrs)** | **Weight (Kg)** | **Clinical Cohort** | **Pathological / Microbiological Findings** |
|---|---|---|---|---|---|
| RM1 | M | 8.28 | 9 | Healthy | N/A |
| RM2 | M | 8.23 | 10.8 | Healthy | N/A |
| RM3 | M | 8.08 | 11.7 | Healthy | N/A |
| RM4 | M | 7.42 | 13.2 | Healthy | N/A |
| RM5 | M | 7.28 | 10.3 | Healthy | N/A |
| **Animal ID** | **Sex** | **Age (yrs)** | **Weight (Kg)** | **Clinical Cohort** | **Pathological / Microbiological Findings** |
| RM6 | M | 7.26 | 12.1 | Healthy | N/A |
| RM7 | M | 7.26 | 11.85 | Healthy | N/A |
| RM8 | M | 6.30 | 10.9 | Healthy | N/A |
| RM9 | M | 6.26 | 10.3 | Healthy | N/A |
| RM10 | M | 6.10 | 11.6 | Healthy | N/A |
| RM11 | M | 5.38 | 11.1 | Healthy | N/A |
| RM12 | M | 5.23 | 11.05 | Healthy | N/A |
| RM13 | M | 8.23 | 9.7 | Healthy | N/A |
| RM14 | M | 5.19 | 10.5 | Healthy | N/A |
| RM15 | M | 5.07 | 10.3 | Healthy | N/A |
| RM16 | F | 11.30 | 7.94 | Healthy | N/A |
| RM17 | F | 8.09 | 7.04 | Healthy | N/A |
| RM18 | M | 6.34 | 10.82 | Healthy | N/A |
| RM19 | M | 5.27 | 8.3 | Healthy | N/A |
| RM20 | M | 13.36 | 6.28 | Diarrhea | Chronic diarrhea with *Campylobacter coli* |
| RM21 | M | 11.19 | 2.9 | Diarrhea | History of fluid diarrhea and cultures of *C. coli* and *Campylobacter jejuni* |
| RM22 | F | 17.34 | 9.67 | Chronic Colitis | History of soft stool and periodic culture of *C. coli.* Generalized hyperplasia and mild inflammation found in stomach and colon |
| RM23 | M | 12.34 | 4.96 | Diarrhea | Mild enteritis with *Balantidium coli* and rare nematodes, 6 days fluid stool |
| **Animal ID** | **Sex** | **Age (yrs)** | **Weight (Kg)** | **Clinical Cohort** | **Pathological / Microbiological Findings** |
| RM24 | M | 11.05 | 2.77 | Diarrhea | History of soft stool due to *C. coli* recovered from gut, inflammatory lesions in stomach and colon |
| RM25 | F | 12.05 | 4.59 | Diarrhea | Chronic recurrent diarrhea with *C. coli* and colitis requiring humane intervention |
| RM26 | F | 12.96 | 3.59 | Diarrhea | Chronic recurrent diarrhea requiring humane intervention, segments of enteritis with cryptitis found in colon with *C. coli* recovered by culture |
| RM27 | F | 17.03 | 9.67 | Chronic Colitis | History of soft stool and chronic colitis and cultures of C. coli and C. jejuni, mild inflammation found in stomach and colon |
| RM28 | F | 8.96 | 8.32 | SIV | Chronically infected SIV w/ daily ART |
| RM29 | F | 7.20 | 6.8 | SIV | Chronically infected SIV w/ daily ART |
| RM30 | F | 6.14 | 7.81 | SIV | Chronically infected SIV w/ daily ART |
| RM31 | M | 6.08 | 11.89 | SIV | Chronically infected SIV w/ daily ART |
| RM32 | M | 5.25 | 9.4 | SIV | Chronically infected SIV w/ daily ART |
| RM33 | M | 6.08 | 11.3 | SIV | Chronically infected SIV w/ daily ART |
| RM34 | M | 5.25 | 7.34 | SIV | Chronically infected SIV w/ daily ART |
| RM35 | M | 5.25 | 8.25 | SIV | Chronically infected SIV w/ daily ART |
| RM36 | M | 4.22 | 5.2 | SIV | Chronically infected SIV w/ daily ART |
| RM37 | M | 4.11 | 6 | SIV | Chronically infected SIV w/ daily ART |

We next examined memory composition changes during chronic diarrheal disease (Fig 5B). Substantial phenotypic remodeling occurred within the Vγ4^+^ subpopulation. Within the colon LPL, Vγ4^+^ T_CM_ frequencies were significantly greater (P = 0.0142; Fig 5B), in contrast from the T_EM_-dominant profile observed under steady-state conditions (Fig 2H). Pie chart compositional analysis further confirmed this phenotypic shift, with peripheral blood Vγ4^+^ T_CM_ cells rising from an average of 7.8% in healthy macaques to 27.11% in diarrhea (Fig 5C) and colonic Vγ4^+^ T_CM_ fraction expanding from 10% in healthy controls to 42.4% during active diarrhea (Fig 5D). This colonic T_CM_ skew resulted in a concomitant contraction of both the Vγ4^+^ T_EM_ fraction (from 66% to 42.6%) and the naive pool (from 24% to 15%). Notably, while the relative proportions shifted robustly within the total Vγ4^+^ population, total absolute frequency changes in individual naive or T_EM_ subsets did not reach statistical significance in either compartment. Conversely, Vδ2^+^ T cells exhibited a significant contraction of the T_CM_ compartment within PBMCs (P < 0.05; Fig 5B), whereas the Vδ1^+^ T cells displayed no significant memory alterations across compartments (Fig 5B, Supplementary Fig 2).

Evaluating transcription factor profiles during chronic inflammation revealed that RORγt expression was significantly and selectively upregulated in colonic Vγ4^+^ T cells during chronic diarrheal disease compared to controls (P < 0.05; Fig 5E), indicating polarization toward a mucosal-protective or pro-inflammatory IL-17-producing phenotype. However, T-bet and EOMES expression remained unaltered across all evaluated subsets in both blood and mucosal compartments (Fig 5F, G).

### Impaired IFN-γ and TNF-α production by circulating Vγ4^+^ T cells during chronic diarrhea

To evaluate effector functions during chronic diarrhea, we measured intracellular cytokines in Vδ1^+^, Vδ2^+^, and Vγ4^+^ T cells following *ex vivo* PMA/ionomycin stimulation of PBMCs (Fig 6). In peripheral blood, chronic diarrhea was associated with impaired Th1 cytokine production by Vγ4^+^ T cells (Fig 6A, B), marked by a significant reduction in IFN-γ compared to healthy controls (P = 0.0043; Fig 6A). Furthermore, TNF-α production was profoundly lost in both circulating Vδ2^+^ (P = 0.0002) and Vγ4^+^ T (P = 0.0001) during active disease (Fig 6B). Although IL-22 and IL-17 frequencies showed downward trends, they did not reach statistical significance due to inter-animal heterogeneity (Fig 6C, D).

**Fig 6.**
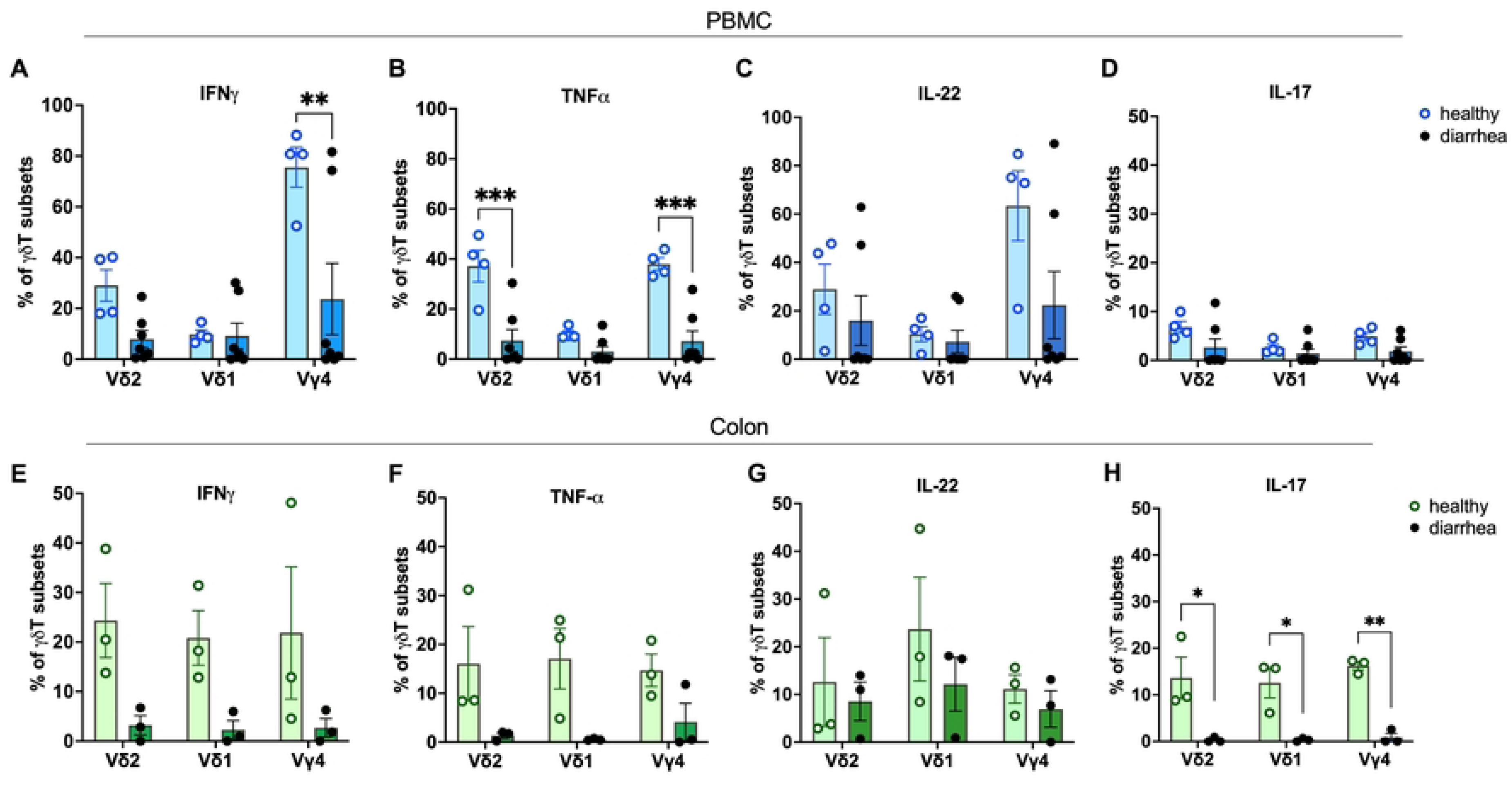
γδ T-cell cytokine responses in PBMC and colon LPL during chronic diarrhea. Frequency of cytokine-producing γδ T-cell subsets (Vδ2+, Vδ1+, and Vγ4+) was quantified by intracellular cytokine staining following ex vivo stimulation with PMA and ionomycin. Comparison of intracellular cytokine expression profiles within PBMCs **(A-D)** and colon LPL **(E-H)** from healthy controls (open symbols and light green/blue bars; n=4–5) and animals with Campylobacter-associated chronic diarrhea (closed symbols and dark green/blue bars; n=3). Panels display the percentage of subsets positive for **(A, E)** IFN-γ, **(B, F)** TNF-α, **(C, G)** IL-22, and **(D, H)** IL-17, tracking simultaneous expression across individual subsets. All values are background-subtracted against unstimulated controls. Data bars represent means ± SEM; each symbol represents an individual animal. Statistical significance was determined using an ordinary two-way ANOVA followed by Tukey’s post-hoc multiple comparisons test; *p<0.05, **p<0.01, ***p<0.001.

We next examined the cytokine functionality locally within the colonic mucosa during chronic diarrheal disease (Fig 6E-H). Despite inter-animal variability, IL-22 emerged as the dominant mucosal response, driven by expanded frequencies in Vδ1^+^ and Vδ2^+^ subsets in two out of the three evaluated animals (Fig 6G). Conversely, IL-17, IFN-γ, and TNF-α response remained low across all subsets. Given that RORγt expression was significantly upregulated in colonic Vγ4^+^ T cells during diarrheal disease (Fig 5E), these data indicate that while chronic intestinal inflammation severely impairs systemic Type 1 effector functions, the local mucosal environment maintains barrier-supportive IL-22 production amid broader Th1/Th17 functional attrition.

### Compartment-specific redistribution and divergent memory remodeling of Vγ4^+^ γδ T cells during chronic SIV infection

Chronic HIV and SIV infections profoundly remodel the γδ T cell compartment(22–26), disrupting tissue-specific dynamics at the host-pathogen interface. Having observed that *Campylobacter*-associated diarrhea promotes a T_CM_-skewed, RORγt^+^ barrier-protective state with maintained IL-22 cytokine producing Vγ4^+^ T cell function in the colon, we investigated whether this barrier-protective program is successfully maintained or disrupted during Antiretroviral Therapy (ART)-suppressed chronic SIV infection. While systemic γδ T cell subsets in PBMCs showed no cross-sectional frequency differences (Fig 7A), or longitudinal alterations (Fig 7B-D), colonic mucosa under effective ART revealed significant expansions of Vδ2^+^ and Vδ1^+^ subsets (P = 0.014 and P = 0.011, respectively), with Vγ4^+^ cells showing an upward trend (Fig 7A). To control for animal-to-animal baseline variation, we leveraged matched longitudinal colon samples, which showed that transition to chronic ART-suppressed SIV infection drove a significant increase in mucosal Vγ4^+^ T cells (P = 0.047; Fig 6G) alongside a strong trend for Vδ1^+^ frequencies (P = 0.079; Fig 7F), whereas Vδ2^+^ subsets remained unchanged. This selective mucosal recovery highlights the distinct homeostatic resilience of the Vγ4/Vδ1 axis during treated viral suppression.

**Fig 7.**
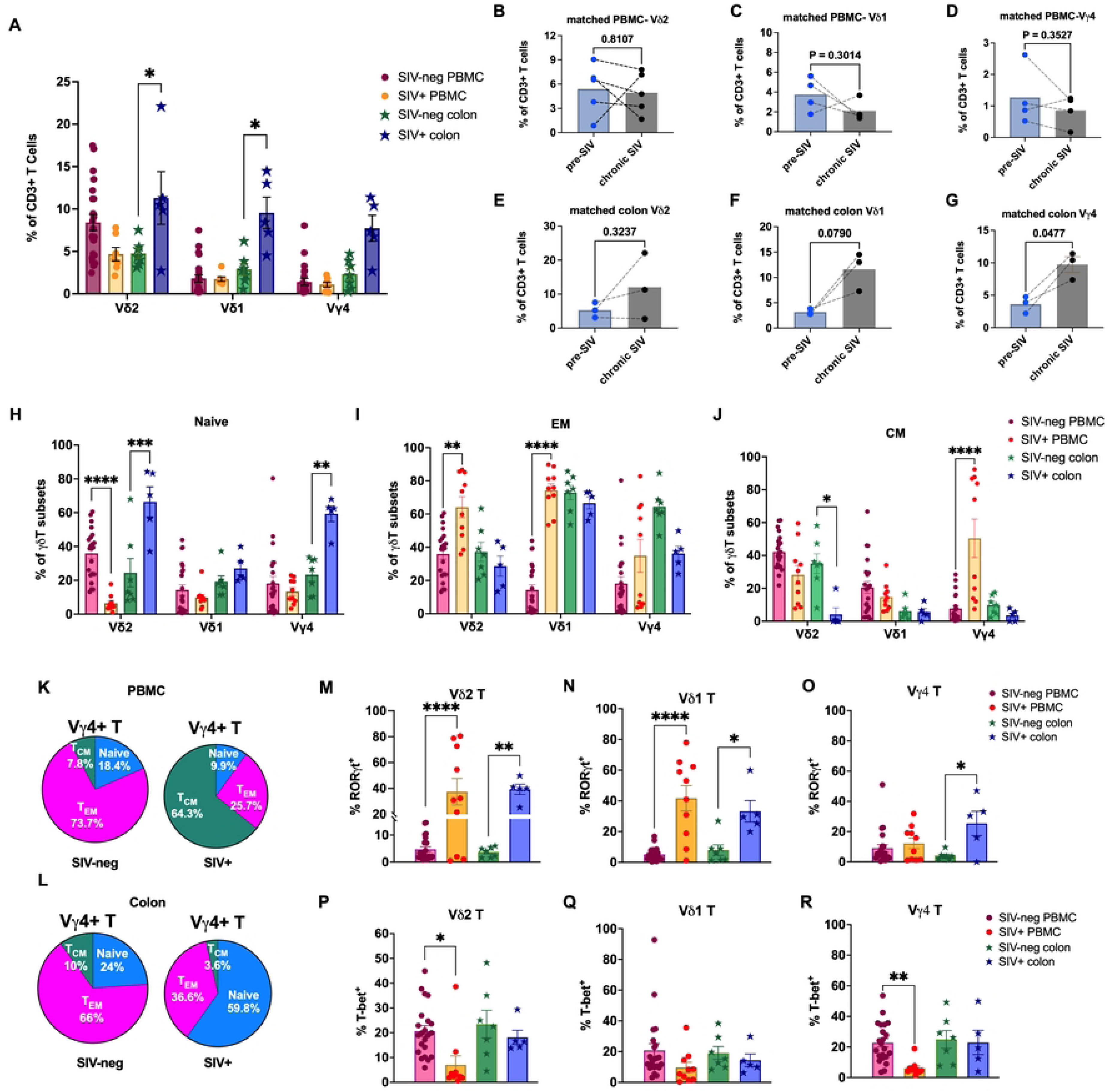
Impact of chronic SIV infection on the frequencies and functional phenotype of macaque γδ T-cell subsets. **(A)** Frequency of Vδ2, Vδ1, and Vγ4 subsets as a percentage of CD3^+^ T cells in PBMCs and colon LPL from animals with chronic SIV infection with ART (PBMC, n=7; colon, n=5) compared to healthy animals (PBMC, n=22; colon, n=7). **(B-G)** Frequencies of each subset in matched samples pre- and post-SIV infection. **(H-J)** Frequencies of CD95^−^CD28^+^ (naive), CD95^+^CD28^−^ effector memory (T_EM_), and CD95^+^CD28^+^ central memory (T_CM_) phenotype cells within Vδ2, Vδ1, and Vγ4 populations in PBMCs and colon LPL from SIV+ and healthy (SIV-neg) animals. Pie charts showing memory subset composition of Vγ4^+^ subsets in PBMCs **(K)** and colon LPL **(L)** between SIV-neg and SIV+ animals. Expression of RORγt **(M-O)**, and T-bet **(P-R)** as a percentage of each γδ subset in PBMCs and colon LPL in SIV-naive versus chronic SIV infection. Bars represent mean ± SEM. Statistical comparisons were performed by two-way ANOVA with Tukey’s multiple comparisons correction. *p<0.05, **p<0.01, ***p<0.001, ****p<0.0001.

Phenotypic profiling revealed divergent systemic and mucosal memory shifts during chronic SIV infection (Fig 7H-L). In PBMCs, Vδ2^+^ and Vδ1^+^ subsets shifted toward terminally differentiated effector phenotypes, marked by significantly contracted circulating naïve Vδ2^+^ pool (p<0.0001; Fig 7H) and expanded T_EM_ fractions (p=0.0017 and p<0.0001, respectively; Fig 7I). Conversely, systemic Vγ4^+^ T cells expanded their central memory (T_CM_) pool from 7.8% in SIV-naïve macaques to 64.3% in the SIV-infected group (p<0.0001; Fig 7J, K). Mucosally, memory dynamics diverged sharply highlighting a unique, tissue-specific disruption across the lineages. Colonic Vδ1^+^ cells remained stable T_EM_ effectors, whereas colonic Vδ2^+^ cells exhibited a significant accumulation of naïve cells (p=0.0001; Fig 7H) with reduced T_CM_ fractions (p=0.0362; Fig 7J). Distinct from chronic diarrhea, mucosal Vγ4^+^ T cells during chronic SIV infection were enriched for naïve subsets (increasing from 24% to 59.8%, p=0.0018; Fig 7H, L), alongside marked contractions in T_EM_ (66% to 36.6%) and T_CM_ (10% to 3.6%) pools.

Transcription factor profiling during chronic SIV infection revealed significantly elevated RoRγt expression in circulating Vδ1^+^ and Vδ2^+^ subsets (p<0.0001, Fig 7M, N) and across all three subsets in colonic LPL (p<0.05, Fig 7M-O). Conversely, T-bet expression was relatively stable in Vδ1^+^ subsets across both compartments (Fig 7Q) but was significantly lower in circulating Vδ2^+^ and Vγ4^+^ cells (p<0.05 and p<0.01 respectively, Fig 7 P, R). No significant changes were observed in EOMES expression (Supplementary Fig 3). Together, these data demonstrate that while chronic SIV infection drives a generalized upregulation of the mucosal barrier-protective factor RORγt, it simultaneously suppresses systemic T-bet expression.

### Significant loss of IFN-γ, TNF-α, and IL-22 effector functions in circulating Vγ4^+^ T cells during chronic SIV infection

We have previously reported that the broader γδ T cell compartment undergoes a profound loss of IL-17 and IL-22 effector functions during chronic, ART-suppressed SIV infection(25). Here, we sought to explore whether these functional deficits are subset-specific (Fig 8). In peripheral blood, chronic SIV infection resulted in a profound, lineage-specific suppression of effector cytokine potency, particularly within the Vδ2^+^ and Vγ4^+^ subsets. Following PMA/ionomycin stimulation of PBMCs, IFNγ production was significantly impaired in circulating Vδ2^+^ (p=0.0004) and Vγ4^+^ (p<0.0001) subsets in chronic SIV-infected macaques compared to healthy controls (Fig 8A). TNFα producing ability was also severely impaired in both Vδ2^+^ and Vγ4^+^ T cells (p<0.0001), whereas it was maintained in the Vδ1^+^ subsets (Fig 8B). Notably, the decline in IL-22 to nearly undetectable levels was highly significant (p<0.0001) in Vγ4^+^ subsets in contrast to their robust production in steady state conditions (Fig 8C). However, systemic IL-17 production remained relatively stable across all three peripheral subsets during chronic SIV infection (Fig 8D), highlighting a lineage-specific functional attrition during chronic SIV infection.

**Fig 8.**
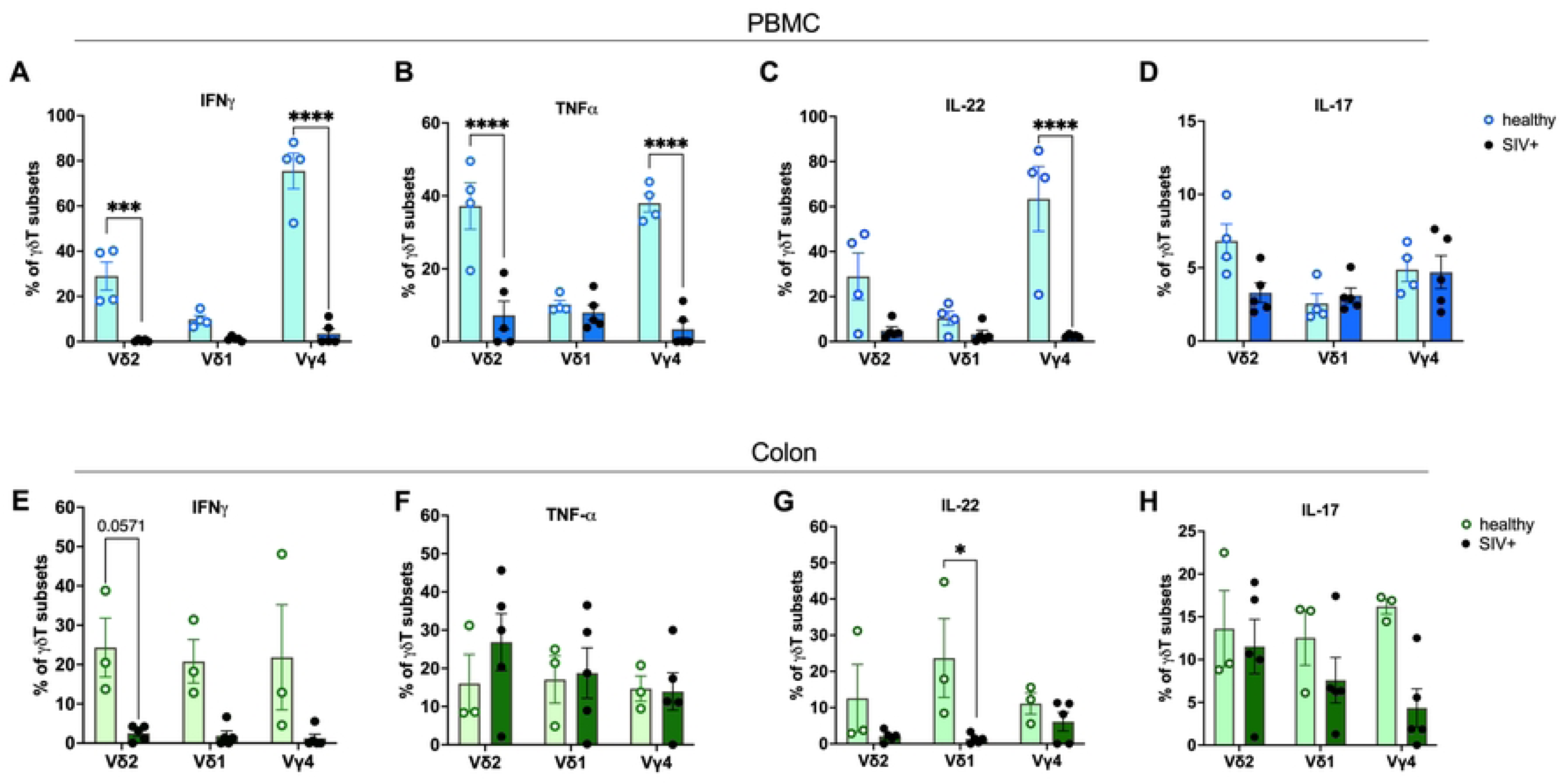
Impact of chronic SIV infection on γδ T-cell cytokine responses in PBMC and colon LPL. Frequency of cytokine-producing γδ T-cell subsets (Vδ2+, Vδ1+, and Vγ4+) was quantified by intracellular cytokine staining following ex vivo stimulation with PMA and ionomycin. Comparison of intracellular cytokine expression profiles within PBMCs **(A-D)** and colon LPL **(E-H)** from healthy controls (open symbols and light green/blue bars; n=3–4) and animals with chronic SIV under ART (closed symbols and dark green/blue bars; n=5). Panels display the percentage of subsets positive for **(A, E)** IFN-γ, **(B, F)** TNF-α, **(C, G)** IL-22, and **(D, H)** IL-17, tracking simultaneous expression across individual subsets. All values are background-subtracted against unstimulated controls. Data bars represent means ± SEM; each symbol represents an individual animal. Statistical significance was determined using an ordinary two-way ANOVA followed by Tukey’s post-hoc multiple comparisons test; *p<0.05, **p<0.01, ***p<0.001.

Within the colonic LPL, the cytokine landscape diverged significantly from systemic circulation (Fig 8E-H). IFNγ production was severely impaired in colonic γδ T cells of macaques with chronic ART-suppressed SIV infection, while trending toward significance in Vδ2^+^ T cells (p=0.0571) due to the small sample size (Fig 8E). Unlike in blood, colonic mucosal TNFα production was preserved across all lineages, maintaining a TNFα-dominant local environment (Fig 8F). Notably, while mucosal IL-22 was significantly depleted within the Vδ1+ subset (p<0.05), it was selectively preserved in colonic Vγ4^+^ subsets (Fig 8G). Mucosal IL-17 production remained stable across all subsets (Fig 8H), demonstrating compartmental preservation of key tissue-protective cytokine programs during chronic ART-suppressed SIV infection. Together, these findings reveal a compartment-specific decoupling of γδ T-cell transcriptional and functional programming during chronic SIV infection. Longitudinally, colonic Vγ4^+^ cells increased, with a trend toward higher Vδ1^+^ cells. Unlike circulating Vδ1^+^ and Vδ2^+^ subsets, which acquired a T_EM_ phenotype, colonic Vγ4^+^ cells exhibited T_CM_ enrichment, suggesting selective homing to the colon followed by AICD. Transcriptional remodeling, marked by elevated systemic RORγt and reduced T-bet was associated with a marked decline in IFNγ and TNFα production in Vδ2^+^ and Vγ4^+^ cells, while IL-17 production was largely preserved.

## Discussion

The present study provides an integrated characterization of a rhesus macaque γδ T-cell population detected using an anti-human Vγ4 antibody and highlights its relevance in mucosal immunity. Complementary TCR sequencing, phenotypic profiling, and functional assays revealed tissue-specific repertoire shifts, including colonic TRGV3 enrichment. The Vγ4 antibody-bound T cells were predominantly Vδ1⁺, displaying an effector-memory phenotype, a T-bet-dominant transcriptional profile, and robust IFN-γ, TNF-α, IL-17, and IL-22 cytokine production. Across two distinct models of chronic intestinal disease (Campylobacter colitis and ART-suppressed SIV), systemic IFN-γ and TNF-α production converged toward significant attrition. Notably, colonic Vγ4⁺ T cells selectively preserved Th17-type responses despite systemic IL-22 loss during chronic SIV infection, demonstrating compartmentalized immune regulation and defining key tissue-specific remodeling dynamics. Together, these findings demonstrate tissue- and disease-associated remodeling of the rhesus Vγ4⁺ γδ T cell subpopulation and provide a foundation for further defining its molecular identity and contribution to mucosal immune regulation.

Our repertoire analyses reveal that γδ T cell organization differs substantially between the γ and δ chains, suggesting distinct evolutionary pressures governing each locus. Uniform *TRG* usage across macaque tissues suggests a conserved structural framework that may be established during thymic development and maintained across peripheral compartments. In contrast, greater *TRD* divergence, particularly expanded splenic *TRD* clonotypes, suggests that δ-chain usage may be more responsive to tissue-specific antigenic or inflammatory cues. This finding aligns with previous observations that Vδ-defined subsets undergo selective peripheral expansion in response to microbial products, stress ligands, and chronic immune activation (27, 28). The dominance of Vδ2 and Vγ9 clonotypes in PBMCs and MLNs is consistent with the well-described prevalence of Vγ9Vδ2 T cells in primate circulation(29, 30), and indicates that MLNs, as gut-draining lymphoid tissues, may serve as trafficking hubs for circulating phosphoantigen-responsive γδ T cells. In contrast, the relative enrichment of Vδ1 usage within the spleen supports growing evidence that Vδ1 populations occupy distinct tissue niches(6, 31, 32) and may undergo localized expansion independent of the dominant Vγ9Vδ2 compartment. Notably, *TRGV3* displayed localized mucosal enrichment in the colonic lamina propria, along with *TRDV1*. While rhesus *TRGV3* shares human *TRGV* sequence similarities, direct equivalence to human *TRGV4* or antibody reactivity remains unconfirmed. Overall, these findings demonstrate that tissue-associated repertoire variation involves subtle shifts in minor V-gene segments alongside clonal expansion, warranting paired sorting and sequencing studies to fully define their molecular identity.

Under homeostatic conditions, rhesus Vγ4⁺ T cells, identified using a cross-reactive human Vγ4 antibody, exhibited features of a highly differentiated tissue-surveillance population. In both blood and colon, Vγ4 preferentially paired with Vδ1, forming a distinct terminal effector-memory cluster separated from Vδ2⁺ T cells, which retained a substantial central memory compartment. These findings parallel recent observations demonstrating that human colonic Vγ4⁺ T cells preferentially associate with Vδ1(18, 33) and exhibit a tissue-adapted phenotype characterized by expression of CD103 and multiple NK receptors, including NKp44, NKG2C, and NKp46(18), supporting a conserved primate mucosal lineage. However, while human studies have primarily captured these cells within specialized tissue niches, their memory dynamics and functional capacity within systemic compartments have remained largely unexplored. Our findings reveal a shared transcriptional landscape across Vγ4⁺, Vδ1⁺, and Vδ2⁺ T cells under steady-state conditions. Systemically and mucosally, Vγ4⁺, Vδ1⁺, and Vδ2⁺ populations shared a Type 1-biased transcriptional program dominated by T-bet and EOMES with minimal RORγt expression. This profile contrasts sharply with murine tissue-resident γδ T cells, which are developmentally programmed toward a RORγt⁺ IL-17-producing phenotype(34, 35), underscoring important species-specific adaptations in primate mucosal immunity.

Functionally, macaque Vγ4⁺ T cells mounted robust polyfunctional cytokine responses following mitogen stimulation, producing IFN-γ, TNF-α, IL-17, and IL-22. This contrasts with human intestinal Vγ2/3/4⁺ cells, which exhibit comparatively reduced IFN-γ and TNF-α production relative to Vδ2⁺ cells following stimulation(18). However, direct comparisons are limited as human studies have largely examined pooled Vγ2/3/4⁺ populations, and the functional profile of purified human Vγ4⁺ T cells remains poorly defined. Together with their transcription factor landscape, this broad effector capacity demonstrates substantial functional plasticity at mucosal barriers. Rather than conforming to a fixed Type 1 or Type 3 lineage, macaque Vγ4⁺ T cells integrate barrier-supportive IL-22 production with a T-bet/EOMES-associated cytotoxic signature, strategically positioning them to couple epithelial maintenance with rapid immune surveillance. Defining these steady-state features provides a framework for understanding how chronic inflammation and viral infection reshape the distribution and function of this γδ T-cell subset across systemic and mucosal compartments.

*Campylobacter*-associated chronic diarrhea induced marked, compartment-specific remodeling of Vγ4⁺ T cells. Circulating Vγ4⁺ T cells exhibited impaired IFN-γ and TNF-α production despite preserved T-bet expression, reflecting systemic Th1 dysfunction. Conversely, colonic Vγ4⁺ T cells expanded their central memory pool, increased RORγt expression, and maintained IL-22 production, indicating local mucosal adaptation to the inflamed intestinal environment. Notably, IL-17 production was profoundly diminished across Vγ4⁺, Vδ1⁺, and Vδ2⁺ subsets. Given the key role of γδ T cell-derived IL-17 in epithelial integrity, antimicrobial peptide expression, recruiting neutrophils, and facilitating early containment of enteric pathogens(36, 37), this broad suppression of innate Th17 responses likely exacerbates barrier breakdown and drives persistent intestinal pathology during chronic enteritis. Consistent with this interpretation, IL-17-producing Vδ1/3⁺ T cells have been associated with enhanced protection against *Clostridioides difficile* infection in children with diarrheal illness(38). The loss of IL-17 across multiple γδ T cell subsets suggests that chronic intestinal inflammation may broadly suppress innate Th17-type responses despite ongoing immune activation. Interestingly, the preservation of colonic Vγ4⁺ IL-22 production together with increased RORγt expression demonstrates selective functional reprogramming rather than global exhaustion. Chronic diarrhea appears to shift γδ T cell responses away from IL-17-mediated inflammatory defense toward an IL-22-dominated tissue-reparative program. Given the critical role of IL-22 in epithelial regeneration, mucus production, and restoration of barrier integrity following injury(39), this shift may serve to limit tissue damage during severe intestinal insult(40). These findings reveal a novel uncoupling of the RORγt-associated IL-17 and IL-22 pathways during chronic diarrheal disease, where tissue-repair mechanisms are preserved despite impaired antimicrobial defense.

Enrichment of phenotypically naïve Vγ4⁺ T cells in the colonic mucosa during chronic SIV infection contrasts sharply with the effector-memory expansion seen in chronic enteropathy, reflecting distinct remodeling mechanisms. Although not directly assessed in this study, it is likely that persistent exposure to viral antigens and translocated microbial products drives chronic activation of mucosal Vγ4⁺ T cells, rendering highly differentiated effector populations susceptible to activation-induced cell death, a mechanism well documented in conventional T cells during HIV and SIV infection(41, 42). While mucosal Vγ4⁺, Vδ1⁺, and Vδ2⁺ subsets upregulated RORγt, indicating broad Type 3 transcriptional polarization, this was not accompanied by preserved effector function. Instead, circulating Vγ4⁺ T cells exhibited marked loss of IFN-γ, TNF-α, and IL-22 alongside reduced T-bet expression. Together, chronic SIV promotes the accumulation of transcriptionally polarized yet functionally impaired γδ T cells, marking progressive mucosal immune dysregulation rather than the tissue-reparative remodeling observed during chronic diarrheal disease.

The shared loss of IFN-γ-producing Vγ4⁺ T cells across chronic diarrhea and SIV infection identifies impairment of mucosal Type 1 immunity as a hallmark of chronic intestinal inflammation. However, while IL-22 production was maintained during chronic enteritis, its severe depletion during chronic SIV infection despite effective viral suppression, highlights distinct outcomes of viral versus non-viral stress.

Because IL-22 directly promotes epithelial regeneration, stem cell renewal, antimicrobial peptide production, and restoration of barrier integrity, loss of IL-22-producing Vγ4⁺ cells may represent a critical mechanism contributing to persistent intestinal epithelial barrier damage during treated HIV/SIV infection. Notably, this functional defect occurred despite effective viral suppression, suggesting that restoration of viral control alone is insufficient to normalize key mucosal immune pathways besides the classical Th17 loss. The loss of IL-17/IL-22 producing cells in the gut during chronic HIV/SIV infection is well established as a key factor for gut barrier disruption and microbial translocation(43–45). Progressive HIV/SIV infection is known to cause the inversion of Vδ1⁺/ Vδ2⁺ frequencies, caused by a preferential expansion of Vδ1⁺ γδ T subsets in the presence of chronic immune activation and microbial translocation(46). The changes observed in Vγ4⁺ T cells likely represent a similar alteration in the mucosal immunity, with loss of barrier protective Type 1 and Type 17 programs beyond the already established Vδ1⁺/ Vδ2⁺ inversion. Furthermore, our earlier studies have demonstrated the selective loss of IL-17/IL-22-producing γδ T cells in the colon during chronic ART-suppressed SIV infection(25, 47). Collectively, these data expand our understanding of mucosal immunity by identifying specific functional shifts within Vγ4^+^ T cells that critically dictate colonic tissue homeostasis during viral infection and inflammatory stress.

Our study has several limitations. The relatively low frequency of the γδ T-cell population detected by the anti-human Vγ4 antibody, particularly in tissue samples, limited sample sizes for some analyses. In addition, repertoire sequencing revealed a trend toward enrichment of rhesus TRGV3 in the colonic lamina propria, although its relationship to the antibody-detected population remains to be established. Sequence comparisons indicate similarities between human TRGV4 and rhesus TRGV gene segments, including TRGV2 and TRGV3, but further analyses are needed to clarify the structural basis of antibody recognition and the contribution of individual rhesus Vγ segments. The observed Vδ1 predominance, effector phenotype, cytokine responses, and colonic enrichment characterize the antibody-detected population and support further investigation of its role in mucosal immunity. Future studies combining TCR sequencing of antibody-sorted cells with single-cell and spatial profiling will help clarify the molecular identity and tissue distribution of this population. Additional investigation of potential interactions with butyrophilin-like molecules, including BTNL3/BTNL8, may also provide insight into the conservation of mechanisms regulating human and rhesus intestinal γδ T cells.

In summary, we present the first characterization of the rhesus macaque Vγ4⁺Vδ1⁺ γδ T cell lineage, defining its specialized roles in mucosal homeostasis. Our data reveal that chronic intestinal inflammation, whether driven by chronic enteritis or SIV infection, induces profound compartment-specific dysregulation within this population. Notably, colonic Vγ4⁺ T cells selectively preserve IL-17 and IL-22 effector capacity during SIV infection, despite significant systemic IL-22 loss, underscoring a key role in local barrier maintenance. Regardless of whether this reflects intrinsic resilience or localized compensation, our data establish Vγ4⁺ T cells as a distinct arm of mucosal immunity with potential therapeutic relevance for HIV-associated enteropathy, inflammatory bowel disease, and chronic intestinal inflammation.

## Materials and Methods

### Animals and sample collection

All animal procedures were approved by the Tulane University Institutional Animal Care and Use Committee (IACUC) and conducted at the Tulane National Biomedical Research Center in strict compliance with NIH guidelines, AAALAC standards, and the Weatherall report recommendations. Rhesus macaques (*Macaca mulatta*) were evaluated across healthy, chronic diarrheal disease, and chronic SIV-infected cohorts. The healthy cohort comprised peripheral blood mononuclear cells (PBMCs; n=22), colon lamina propria lymphocytes (LPLs; n=7), liver cells (n=3), and pooled duodenal and colonic LPLs (n=3). The diarrheal disease cohort comprised seven Campylobacter-associated cases, six with chronic diarrhea and one with chronic colitis, with colon LPLs analyzed from four animals after excluding one sample due to low viability. Animals in this cohort were euthanized due to severe clinical disease based on veterinary assessment (Table 1). The chronic SIV-infected cohort included animals contributing PBMCs (n=10) and colon LPLs (n=5). Macaques were infected intrarectally with SIVmac_251_ and sampled during chronic infection at least four months post-infection. All SIV-infected animals received combination antiretroviral therapy (cART) for at least three months, consisting of daily subcutaneous tenofovir disoproxil fumarate (5.1 mg/kg), emtricitabine (30 mg/kg), and dolutegravir (2.5 mg/kg).

### Tissue processing and Cell Isolation

Peripheral blood collected in EDTA tubes was processed immediately by density gradient centrifugation over Lymphocyte Separation Medium at 1500 rpm for 45 minutes to isolate PBMCs. Spleen and liver tissues were manually disrupted in cold PBS with 2% fetal calf serum using a 16-gauge needle, passed through a 100-µm cell strainer, and centrifuged at 600 × g for 7 minutes. Ammonium-Chloride-Potassium (ACK) lysis was performed as necessary for red blood cell removal. Colon LPLs were isolated using Percoll density gradient centrifugation(47), washed, and resuspended in complete media. Cell viability across all preparations was confirmed by trypan blue exclusion, and isolated cells were resuspended at 1 × 10^6^ cells/mL in complete media for downstream phenotypic and functional assays.

### Immune phenotyping and functional assessment

Multi-color flowcytometric analysis was performed on cells according to standard procedures using anti-human mAbs that cross-react with rhesus macaques (Table 2). Surface staining for CD45, CD3, TCR γδ, TCR Vγ4, TCR Vδ1, TCR Vδ2, CD4, CD8, CD95, CD28, CD14, PD-1, CD20, α4β7, and CD127 was performed at 4°C for 30 min. as earlier described(47). Intracellular transcription factors (RoRγt, T-bet, and EOMES) were stained following fixation and permeabilization using the True-Nuclear Transcription Factor Buffer Set (BioLegend) according to the manufacturer’s protocol. To determine functional capacity, cells were stimulated with either phorbol 12-myristate 13-acetate (PMA; 50 ng/mL; Sigma-Aldrich) and ionomycin (1 µg/mL; ThermoFisher Scientific) or anti-CD3/CD28 antibodies (5 µg/mL; NHP Reagent Resource). Brefeldin A (1 µg/mL; BD Biosciences) was added 1 h post-stimulation, and cells were incubated for 16 h at 37°C in 5% CO_2_ incubator. Following surface staining, cells were fixed and permeabilized using the Cytofix/Cytoperm kit (BD Biosciences) and stained for intracellular cytokines (IL-17, IL-22, TNF-α, and IFN-γ). Samples were acquired on a BD FACSymphony A5 flow cytometer (BD Biosciences) capturing 200,000 lymphocyte events per sample. Data were analyzed using FlowJo software (v10; BD Biosciences), and background subtraction using unstimulated control values was applied to calculate net cytokine responses.

**Table 2.**
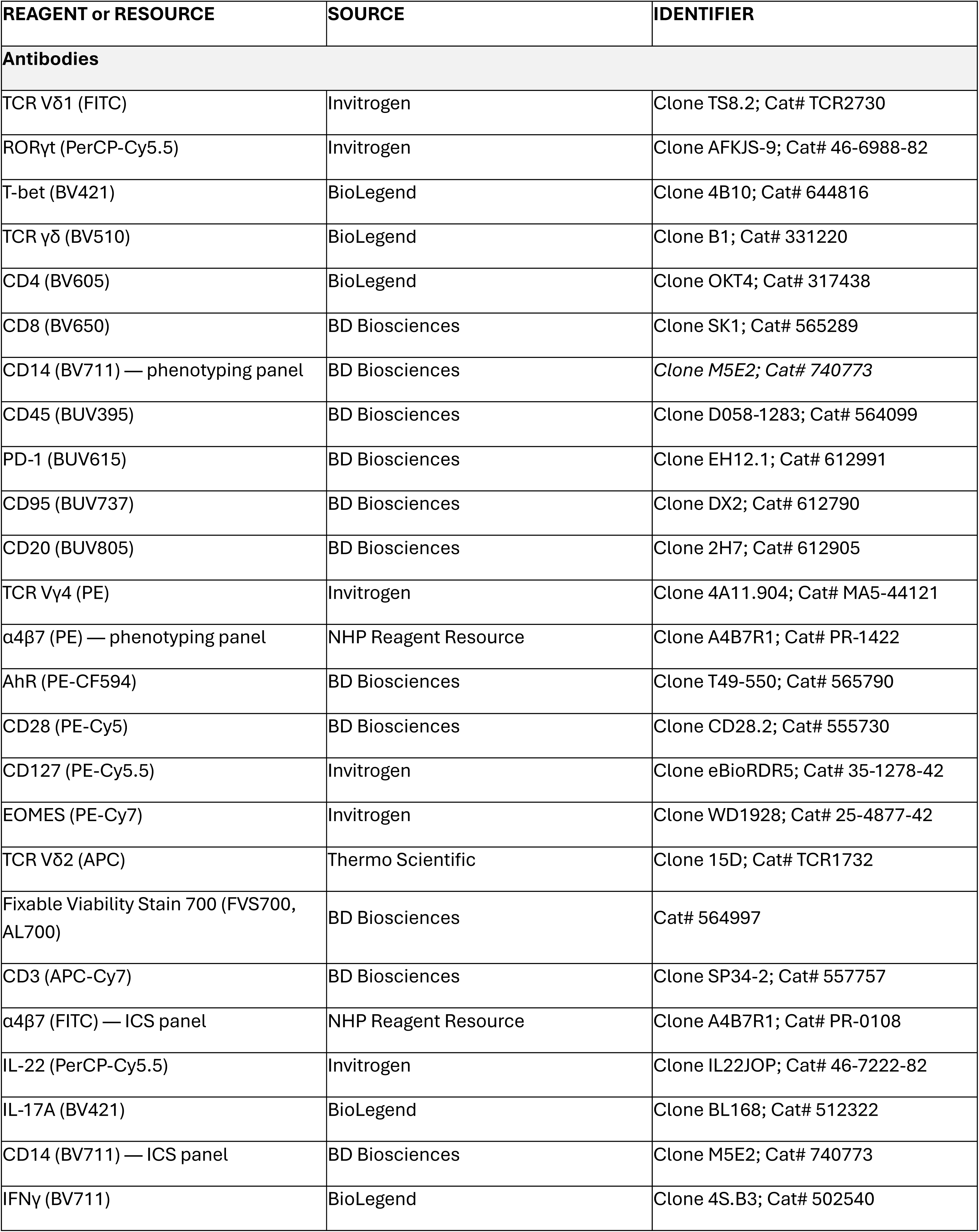
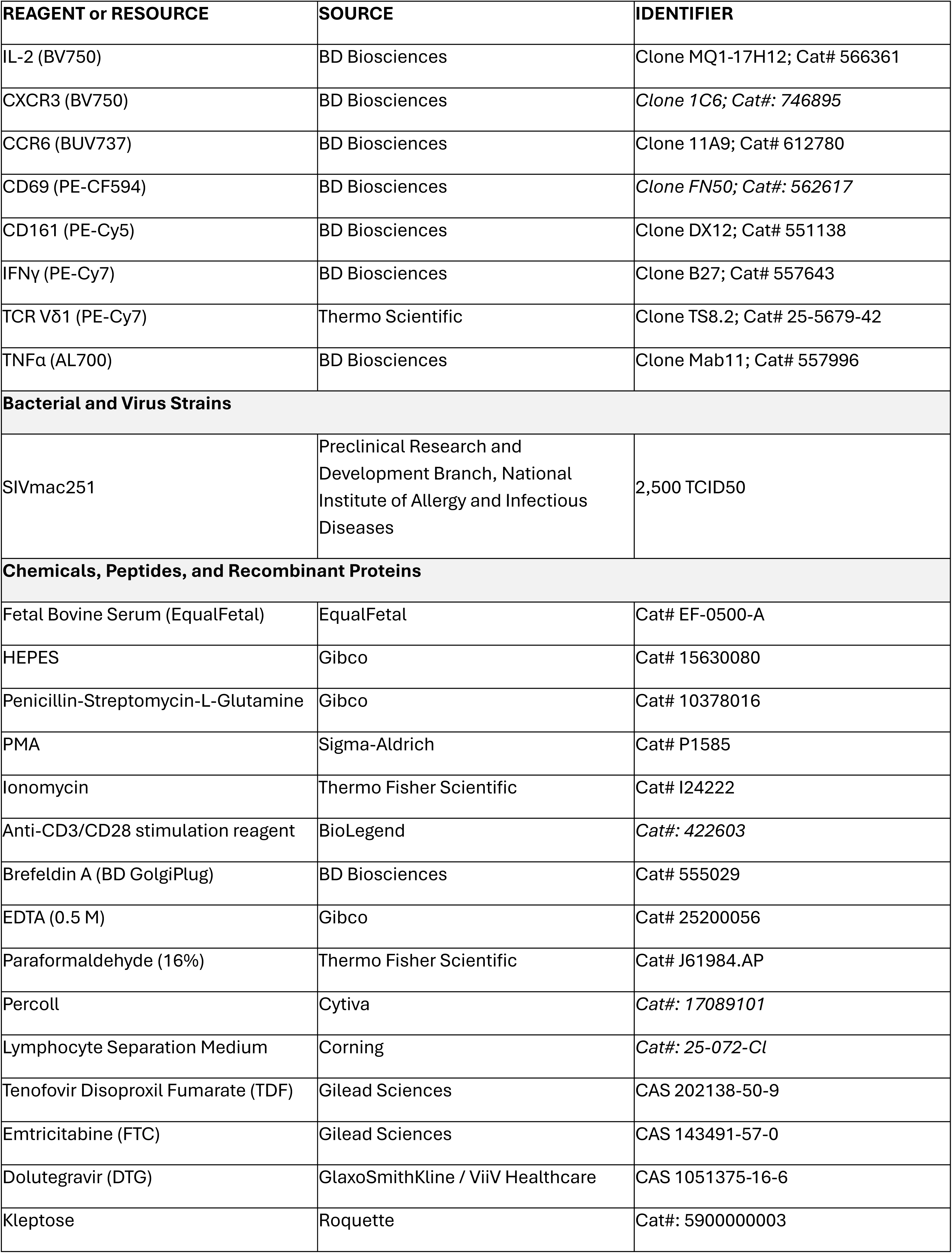
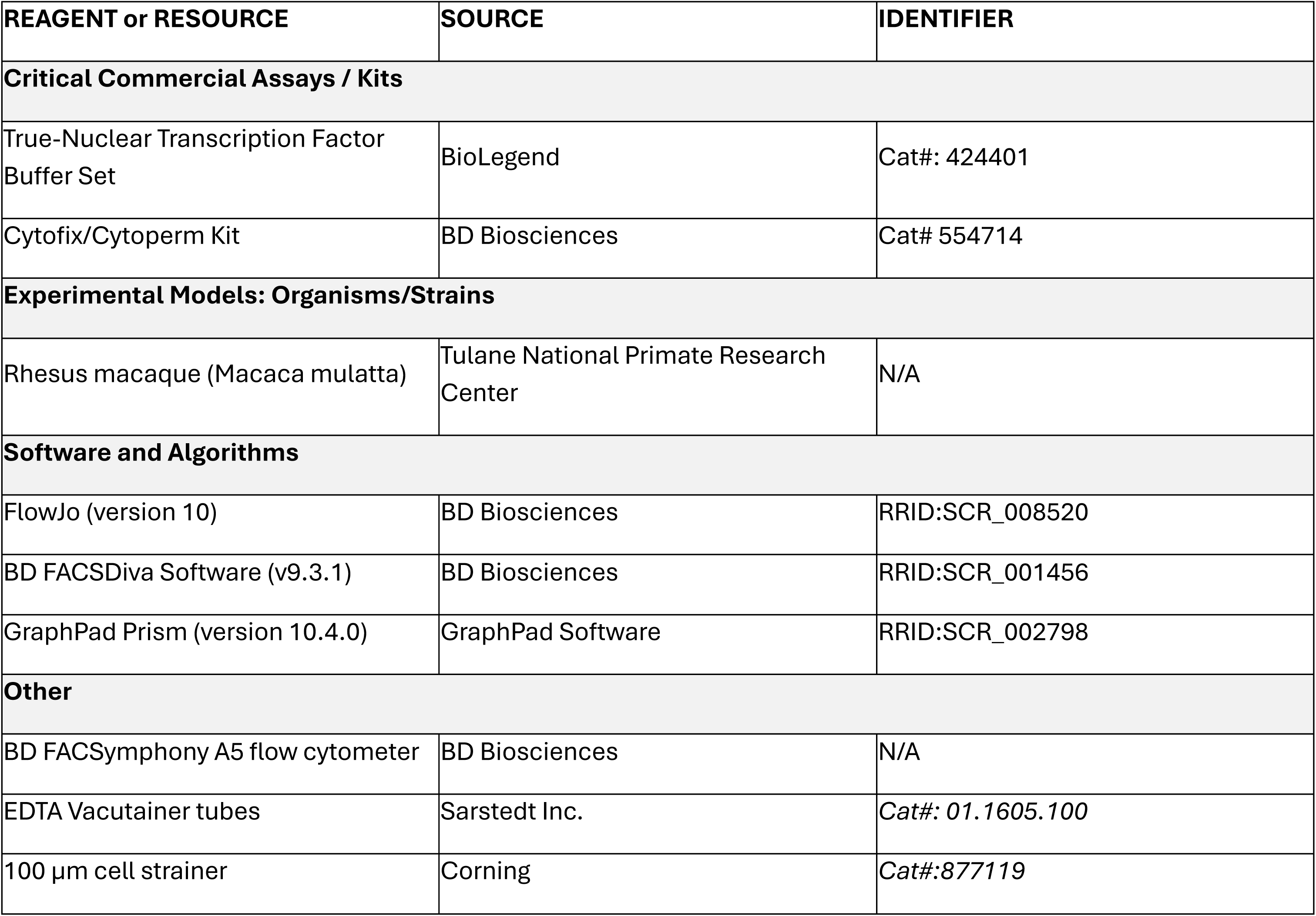
Key resources.

| **REAGENT or RESOURCE** | **SOURCE** | **IDENTIFIER** |
|---|---|---|
| **Antibodies** |  |  |
| TCR Vδ1 (FITC) | Invitrogen | Clone TS8.2; Cat# TCR2730 |
| RORγt (PerCP-Cy5.5) | Invitrogen | Clone AFKJS-9; Cat# 46-6988-82 |
| T-bet (BV421) | BioLegend | Clone 4B10; Cat# 644816 |
| TCR γδ (BV510) | BioLegend | Clone B1; Cat# 331220 |
| CD4 (BV605) | BioLegend | Clone OKT4; Cat# 317438 |
| CD8 (BV650) | BD Biosciences | Clone SK1; Cat# 565289 |
| CD14 (BV711) — phenotyping panel | BD Biosciences | *Clone M5E2; Cat# 740773* |
| CD45 (BUV395) | BD Biosciences | Clone D058-1283; Cat# 564099 |
| PD-1 (BUV615) | BD Biosciences | Clone EH12.1; Cat# 612991 |
| CD95 (BUV737) | BD Biosciences | Clone DX2; Cat# 612790 |
| CD20 (BUV805) | BD Biosciences | Clone 2H7; Cat# 612905 |
| TCR Vγ4 (PE) | Invitrogen | Clone 4A11.904; Cat# MA5-44121 |
| α4β7 (PE) — phenotyping panel | NHP Reagent Resource | Clone A4B7R1; Cat# PR-1422 |
| AhR (PE-CF594) | BD Biosciences | Clone T49-550; Cat# 565790 |
| CD28 (PE-Cy5) | BD Biosciences | Clone CD28.2; Cat# 555730 |
| CD127 (PE-Cy5.5) | Invitrogen | Clone eBioRDR5; Cat# 35-1278-42 |
| EOMES (PE-Cy7) | Invitrogen | Clone WD1928; Cat# 25-4877-42 |
| TCR Vδ2 (APC) | Thermo Scientific | Clone 15D; Cat# TCR1732 |
| Fixable Viability Stain 700 (FVS700, AL700) | BD Biosciences | Cat# 564997 |
| CD3 (APC-Cy7) | BD Biosciences | Clone SP34-2; Cat# 557757 |
| α4β7 (FITC) — ICS panel | NHP Reagent Resource | Clone A4B7R1; Cat# PR-0108 |
| IL-22 (PerCP-Cy5.5) | Invitrogen | Clone IL22JOP; Cat# 46-7222-82 |
| IL-17A (BV421) | BioLegend | Clone BL168; Cat# 512322 |
| CD14 (BV711) — ICS panel | BD Biosciences | Clone M5E2; Cat# 740773 |
| IFNγ (BV711) | BioLegend | Clone 4S.B3; Cat# 502540 |
| **REAGENT or RESOURCE** | **SOURCE** | **IDENTIFIER** |
| IL-2 (BV750) | BD Biosciences | Clone MQ1-17H12; Cat# 566361 |
| CXCR3 (BV750) | BD Biosciences | *Clone 1C6; Cat#: 746895* |
| CCR6 (BUV737) | BD Biosciences | Clone 11A9; Cat# 612780 |
| CD69 (PE-CF594) | BD Biosciences | *Clone FN50; Cat#: 562617* |
| CD161 (PE-Cy5) | BD Biosciences | Clone DX12; Cat# 551138 |
| IFNγ (PE-Cy7) | BD Biosciences | Clone B27; Cat# 557643 |
| TCR Vδ1 (PE-Cy7) | Thermo Scientific | Clone TS8.2; Cat# 25-5679-42 |
| TNFα (AL700) | BD Biosciences | Clone Mab11; Cat# 557996 |
| **Bacterial and Virus Strains** |  |  |
| SIVmac251 | Preclinical Research and Development Branch, National Institute of Allergy and Infectious Diseases | 2,500 TCID50 |
| **Chemicals, Peptides, and Recombinant Proteins** |  |  |
| Fetal Bovine Serum (EqualFetal) | EqualFetal | Cat# EF-0500-A |
| HEPES | Gibco | Cat# 15630080 |
| Penicillin-Streptomycin-L-Glutamine | Gibco | Cat# 10378016 |
| PMA | Sigma-Aldrich | Cat# P1585 |
| Ionomycin | Thermo Fisher Scientific | Cat# I24222 |
| Anti-CD3/CD28 stimulation reagent | BioLegend | *Cat#: 422603* |
| Brefeldin A (BD GolgiPlug) | BD Biosciences | Cat# 555029 |
| EDTA (0.5 M) | Gibco | Cat# 25200056 |
| Paraformaldehyde (16%) | Thermo Fisher Scientific | Cat# J61984.AP |
| Percoll | Cytiva | *Cat#: 17089101* |
| Lymphocyte Separation Medium | Corning | *Cat#: 25-072-Cl* |
| Tenofovir Disoproxil Fumarate (TDF) | Gilead Sciences | CAS 202138-50-9 |
| Emtricitabine (FTC) | Gilead Sciences | CAS 143491-57-0 |
| Dolutegravir (DTG) | GlaxoSmithKline / ViiV Healthcare | CAS 1051375-16-6 |
| Kleptose | Roquette | Cat#: 5900000003 |
| **REAGENT or RESOURCE** | **SOURCE** | **IDENTIFIER** |
| **Critical Commercial Assays / Kits** |  |  |
| True-Nuclear Transcription Factor Buffer Set | BioLegend | Cat#: 424401 |
| Cytofix/Cytoperm Kit | BD Biosciences | Cat# 554714 |
| **Experimental Models: Organisms/Strains** |  |  |
| Rhesus macaque (Macaca mulatta) | Tulane National Primate Research Center | N/A |
| **Software and Algorithms** |  |  |
| FlowJo (version 10) | BD Biosciences | RRID:SCR_008520 |
| BD FACSDiva Software (v9.3.1) | BD Biosciences | RRID:SCR_001456 |
| GraphPad Prism (version 10.4.0) | GraphPad Software | RRID:SCR_002798 |
| **Other** |  |  |
| BD FACSymphony A5 flow cytometer | BD Biosciences | N/A |
| EDTA Vacutainer tubes | Sarstedt Inc. | *Cat#: 01.1605.100* |
| 100 µm cell strainer | Corning | *Cat#:877119* |

### TCR Sequence Analysis

αβTCR-negative T cells were lysed in RLT Plus buffer (Qiagen) supplemented with 1% β-mercaptoethanol (β-ME). Genomic DNA was isolated using the AllPrep DNA/RNA Micro Kit (Qiagen, cat# 80284). Multiplex PCR amplification of TCR gamma (TRG) and delta (TRD) V–J junctions was performed using Platinum Taq DNA Polymerase (Invitrogen, cat#10966026) with specific forward V-region and reverse J-region primers. The TRG primer mix comprised the following forward V-region primers: TRGV1, 5′-AGTATGACACTGCAAGCACAAGGAA-3′; TRGV2, 5′-CTTATGGAAGCACAAGGAGCAA-3′; TRGV3, 5′-TATACTCATACACGCACAAGGTGG-3′; TRGV8, 5′-ACAACTTGAGATTGAGACTGCGAA-3′; TRGV9, 5′-CCATCGGGCAAATTTGAGGT-3′; and TRGV10, 5′-ACCGAATCAGGCTTTGGAGC-3′. Reverse J-region primers were TRGJ1-1, 5′-CAGCAAATATCTTGATCCAACCAG-3′; TRGJ1-2, 5′-TGTTCCGGGACCAAATAGCTTG-3′; TRGJ2-1, 5′-TGAGCCTAGTCCCTTCAGCA-3′; and TRGJ2-3, 5′-ACAAGTGTTGTTCCACTGCCA-3′. The TRD primer mix comprised the following forward V-region primers: TRDV1-1, 5′-AAGCAGCTAGCTCCATCACCG-3′; TRDV1, 5′-TGGTCGCTATTCTGTCAACTTCA-3′; TRDV4, 5′-CAGGAGGCGACCTGAAGAAC-3′; TRDV2, 5′-GGCCCTGGTTTCAAGGACAA-3′; and TRDV3, 5′-ACGGTTTTCTGTGAAGCACC-3′. Reverse J-region primers were TRDJ1, 5′-TTCCACAGTCACACGGGTTC-3′; TRDJ4, 5′-CCAAAGATCAGTGGTCTGGCA-3′; TRDJ2, 5′-TCCACGATGAGTTGAGTTCCC-3′; and TRDJ3, 5′-AAACATCTGTCGGGTGTCCC-3′.

Amplified products (100–300 bp) were gel-purified from 2% agarose gels using the NucleoSpin Gel and PCR Clean-up kit (Macherey-Nagel, cat#740609.250) and subjected to Illumina sequencing. Adaptive Immune Receptor Repertoire (AIRR) formatted sequencing files were filtered to remove singletons to minimize sequencing artifacts and analyzed using the immunarch package in R [ImmunoMind Team. (2019). immunarch: An R Package for Painless Bioinformatics Analysis of T-Cell and B-Cell Immune Repertoires. Zenodo. http://doi.org/10.5281/zenodo.3367200].

### Statistical analyses

Statistical analysis was performed using GraphPad Prism software (v10.4.0). Multi-group comparisons were evaluated using two-way analysis of variance (ANOVA) followed by Tukey’s post-hoc test for multiple comparisons. Correlations between cellular frequencies were assessed using Spearman rank correlation. Paired healthy and SIV-infected timepoint samples (n = 4) were analyzed using paired two-tailed t-tests. Data are presented as mean ± standard error of the mean (SEM), with statistical significance defined as p < 0.05.

## Acknowledgements

We thank the Tulane National Biomedical Research Center (TNBRC; RRID: SCR_008167), its veterinary staff, and the Flow Cytometry Core (S10 OD026800, RRID: SCR_024611/SCR_008167). We acknowledge ViiV and Gilead for antiretroviral drugs.

## Data availability

The TCRSeq datasets generated and analyzed during the current study have been deposited in the NCBI Gene Expression Omnibus (GEO) under accession number GSE352564. Other data available from this study are available from the corresponding author upon reasonable request.

## Author contributions

**ABM:** Writing – original draft, Methodology, Investigation, Formal analysis, Data curation. **ARR:** Writing – review & editing, Methodology, Investigation, Formal analysis. **TEM:** Methodology, Investigation. **VAH:** Methodology, Investigation. **JD:** Methodology, Investigation. **ST:** Writing – review & editing. **YF:** Writing – review & editing. **NB:** Writing – review & editing. **JMB:** Writing – review & editing, Investigation, Supervision. **NR:** Writing – original draft, review & editing, Supervision, Investigation, Funding acquisition, Conceptualization.

## Supporting information captions

**Supplementary Fig 1. Co-expression of Vγ4^+^ TCR with Vδ1^+^ and Vδ2^+^ TCR in macaque colonic lamina propria cells. (A)** Representative co-expression dot plot and correlation between Vγ4 and Vδ1 T cell frequencies in colon LPL (n=7) showing significant positive correlation (Spearman r=0.8571, p=0.0238). **(B)** Representative co-expression dot plot and correlation between Vγ4 and Vδ2 frequency in the same LPL samples showing no correlation.

**Supplementary Fig 2. Naïve and effector memory phenotype and transcription factor expression by circulating and colonic γδ T subsets in healthy and diarrhea-inflicted macaques. (A)** Frequency of CD95^−^CD28^+^ naïve and **(B)** CD95^+^CD28^−^ effector memory (T_EM_) phenotype cells within Vδ2, Vδ1, and Vγ4 populations in PBMCs and colon LPL from animals with chronic diarrheal disease (PBMC, n=7; colon LPL, n=4) compared to healthy animals (PBMC, n=22; colon LPL, n=7). **(C)** Expression of EOMES as a percentage of each γδ T subset in PBMCs and colon LPL in healthy versus chronic diarrheal disease. Bars represent mean ± SEM. No significant differences were found. Statistical comparisons were performed by two-way ANOVA with Tukey’s multiple comparisons correction.

**Supplementary Fig 3. EOMES expression by circulating and colonic γδ T subsets from healthy and chronic SIV-infected macaques.** Percentage of EOMES^+^ cells within each γδ T subset in PBMCs and colon LPL from healthy SIV-naïve versus SIV^+^ macaques. Bars represent mean ± SEM. No significant differences were found. Statistical comparisons were performed by two-way ANOVA with Tukey’s multiple comparisons correction.

